# Spatiotemporally restricted Hippo signalings instruct the fate and maturation of hepatobiliary cells

**DOI:** 10.1101/2024.11.02.621695

**Authors:** Yebin Wang, Zhenxing Zhong, Yu Wang, Ying Zhu, Kun-Liang Guan, Fa-Xing Yu

## Abstract

The Hippo pathway is an evolutionary conserved signaling cascade involved in organ size control and tumorigenesis, but its cellular functions and regulatory mechanisms are not fully understood. Recently, we have defined two independent modules (HPO1 and HPO2) in the Hippo signaling network. Here, by using spatially resolved transcriptomic and imaging analysis of mouse livers with defective Hippo signaling, we show that HPO1 and HPO2 operate in distinct cells at different developmental stages to regulate the fate and maturation of liver parenchymal cells. HPO1 controls the maturation of hepatocytes postnatally, and its perturbation leads to the expansion of immature hepatocytes (imHep). HPO2, on the other hand, regulates the maturation of cholangiocytes perinatally, and its ablation results in the accumulation of immature cholangiocytes (imCho2) identical to developing ductal plate cells. Moreover, the inactivation of HPO1 or HPO2 causes the conversion of hepatocytes into immature cholangiocytes (imCho1). These immature cells are also observed in regenerating livers following different damages. In contrast, deletion of *Yap/Taz* encoding downstream effectors of the Hippo pathway accelerates liver maturation and promotes cell death. These findings suggest that the spatiotemporally restricted Hippo signaling modules act as checkpoints in liver development and may coordinate cell proliferation and maturation to ensure proper liver size and function.

## INTRODUCTION

The liver is a central metabolic organ that frequently encounters tissue damage caused by drugs or infections, and persistent tissue damages lead to liver fibrosis, cirrhosis, and tumorigenesis. Currently, chronic liver disease and liver cancer are major causes of human death ^1-3^. The liver parenchymal contains hepatocytes and cholangiocytes (biliary epithelial cells); both cell types (together called hepatobiliary cells) are specified from hepatoblasts (bipotential progenitors) in embryonic development, and after birth, hepatobiliary cells continue to develop until a functional matured liver is established ^4-7^. Hepatobiliary cells may dedifferentiate, transdifferentiate, and proliferate during liver regeneration to repopulate liver parenchyma ^8^. However, it is currently unclear how different signals orchestrate cell fate, plasticity, proliferation, and functional maturation to regulate liver development, homeostasis, regeneration, and disease progression.

The Hippo pathway has been established as a signaling cascade regulating tissue development and organ size ^9-13^. In the Hippo pathway, downstream effectors YAP and TAZ are repressed by multiple upstream kinases (MST1/2, MAP4K1-7, and LATS1/2) and scaffolding proteins (SAV1, NF2, MOB1, and WWC1-3). YAP/TAZ, as transcriptional co-factors, modulates the expression of many genes involved in cell proliferation and survival, hence determining organ size by regulating cell numbers ^9-12^. In mouse livers, transgenic expression of *Yap* or genetic deletion of upstream negative regulators results in a robust increase in liver size and tumorigenesis ^14-20^. However, the cell proliferation and liver size do not change significantly upon conditional deletion of *Yap* and *Taz* ^21-24^. Moreover, recent findings suggest that the liver enlargement induced by YAP hyperactivation deviates from developmental growth ^22^. These counterintuitive results have challenged the prevailing view of Hippo signaling as a master organ size regulator. Hence, the Hippo pathway may not simply count and control cell numbers during development, and it is important to explore its upstream signals and genuine cellular functions.

Recently, we have revealed that HPO1 (MST1/2–SAV1–WWC1-3–LATS1/2) and HPO2 (MAP4K1-7–NF2–LATS1/2), two largely independent signaling modules, jointly regulate the activity of LATS1/2 and YAP/TAZ, and genetic perturbations of HPO1 and/or HPO2 genes in mouse livers result in drastically different phenotypes, such as liver sizes and tumor subtypes (Fig 1A) ^21,25,26^. In this study, by using single-cell RNA sequencing (scRNA-seq), spatial transcriptomics (ST), imaging, and mouse genetics, we show that HPO1 predominantly operates in hepatocytes postnatally, whereas HPO2 mainly functions in cholangiocytes perinatally, and deletion of HPO1 and/or HPO2 genes results in three pools of cells resembling immature hepatobiliary cells at different developmental or regenerating stages. In contrast, deletion of *Yap/Taz* speeds up liver maturation. We propose that HPO1 and HPO2 serve as maturation checkpoints for hepatocytes and cholangiocytes, respectively, and may coordinate cell proliferation and cell maturation to determine liver growth, size, and function.

**Figure 1.**
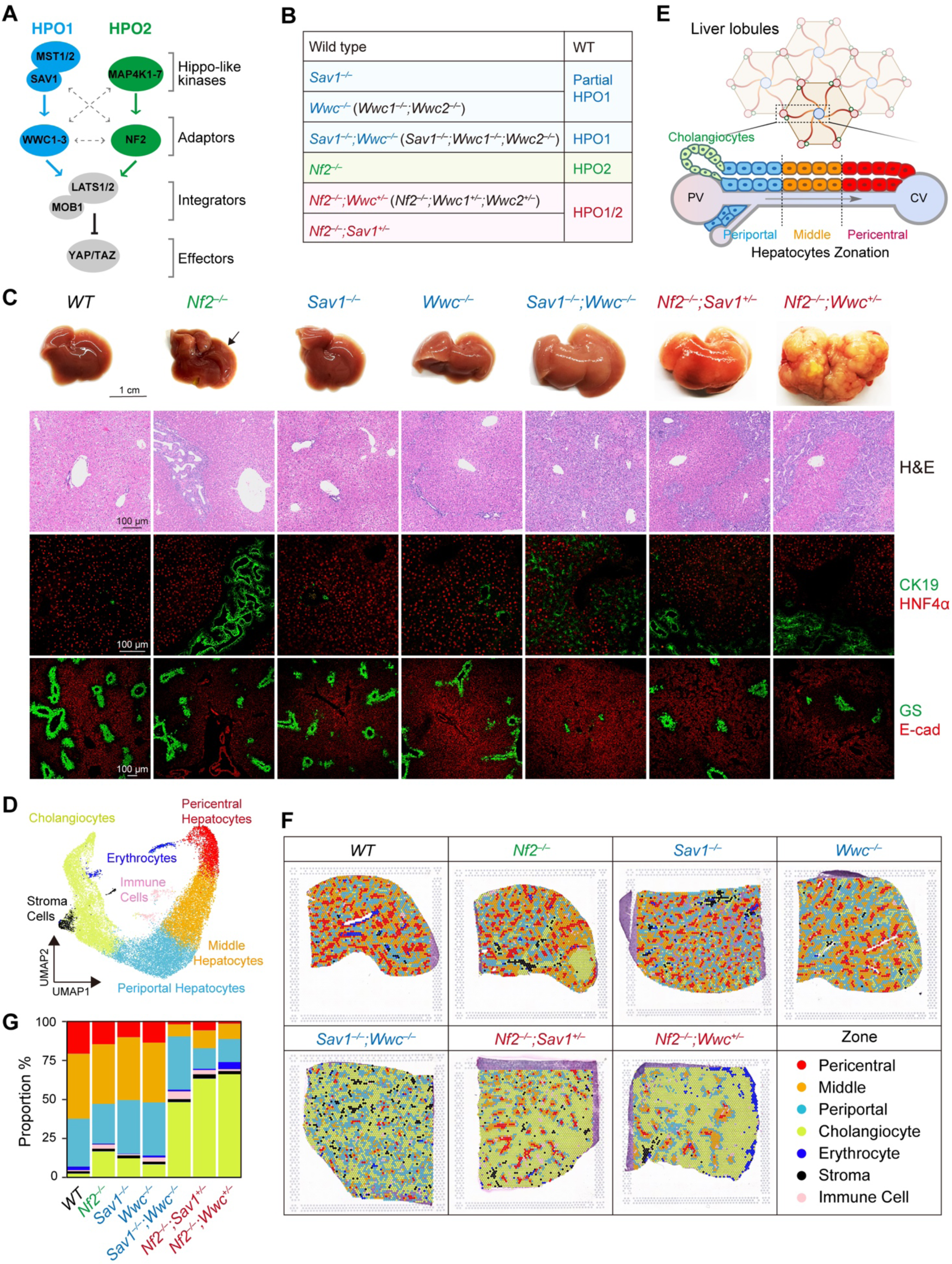
Hippo signaling inactivation results in diverse liver developmental defects. (A) Diagram illustrating two Hippo signaling modules. Upstream regulators of YAP/TAZ are classified into two Hippo signaling modules: HPO1 and HPO2, each of which can independently regulate YAP/TAZ activity. (B) Table summarizing mouse models used in this study, genotypes contain HPO1 and/or HPO2 deletions. (C) Deletion of Hippo pathway genes results in distinct liver morphologies. Gross liver images, H&E CK19, HNF4α, GS, and Ecad staining are shown. 3-week-old wild type (WT) mice or mice with liver-specific knockout of different HPO modules. Scale bars: 1 cm or 100 μm as indicated. (D) Integrated and annotated UMAP embedding of spatial spots. The spatial spots are divided into 5 clusters based on their molecular signatures, and the hepatocyte zone can be further subdivided into three sub-clusters based on its zonation signature. (E) Diagram illustrating liver zonation. The liver consists of repeating units called liver lobules, which are highly organized structures. The liver lobule can be divided into three zones: the pericentral zone, the middle zone, and the periportal zone. (F) Spatial projection of cell clusters onto H&E staining images of indicated livers. (G) Proportions of annotated zones in livers with different genotypes.

## RESULTS

### ST analysis reveals dramatic changes in livers with defective Hippo signaling

In the Hippo signaling network, WWC1/2 and SAV1 mainly mediate LATS1/2 activation by MST1/2 (HPO1), and NF2 mainly mediates LATS1/2 activation by MAP4K1-7 (HPO2). Hence, as adaptors, NF2, WWC1-3, and SAV1 transmit signals from Hippo-like kinases to LATS1/2 and YAP/TAZ (Fig 1A)^25^. We generated different mouse strains with liver-specific deletion (*Albumin*-cre) of single or multiple genes resembling HPO1 and/or HPO2 inactivation (Fig 1B). For simplicity, *Wwc^+/–^* and *Wwc^−/−^* were used throughout this study for *Wwc1^+/–^;Wwc2^+/–^*and *Wwc1^−/−^;Wwc2^−/−^* genotypes (no *Wwc3* gene in mice), respectively. During postnatal mouse development, 21-day-old (P21) appeared to be a critical time point for liver growth, as the buildup of liver mass started to decelerate (Fig S1A-C). On P21, livers with both HPO1 and HPO2 inactivation (*Nf2^−/−^;Wwc^+/–^*or *Nf2^−/−^;Sav1^+/–^*) or complete HPO1 inactivation (*Sav1^−/−^;Wwc^−/−^*) were dramatically enlarged, whereas livers with HPO2 inactivation (*Nf2^−/−^*) or partial HPO1 inactivation (*Sav1^−/−^* or *Wwc^−/−^*) were not enlarged robustly (Fig 1C). *Nf2^−/−^* livers frequently developed bile duct hamartomas in peripheral regions, consistent with previous reports (Fig 1C) ^17^. *Nf2^−/−^;Wwc^−/−^* or *Nf2^−/−^;Sav1^−/−^*mice died within a week after birth and were not included in this study ^25^. Hepatocytes and cholangiocytes are marked by the expression of HNF4α and CK19, respectively. Cholangiocytes were expanded in *Nf2^−/−^* livers, further increased in *Nf2^−/−^;Wwc^+/–^*, *Nf2^−/−^;Sav1^+/–^*, and *Sav1^−/−^;Wwc^−/−^* livers, whereas the increase in *Sav1^−/−^* or *Wwc^−/−^* livers were less robust (Fig 1C). These results indicate that the inactivation of HPO1 and/or HPO2 affects liver size and hepatobiliary differentiation differently. A comprehensive analysis of liver cells at a higher resolution in space and time is required to understand dynamic changes in cell types and states in these livers.

The explosive development of ST technologies provides a unique opportunity to resolve complex tissue *in situ* ^27,28^. To survey the changes in cellular compositions associated with the inactivation of Hippo signalings, we perform ST analysis (Visium) for tissue sections from livers with genetic deletion of HPO1 and/or HPO2 genes. After filtering out the low-quality spatial spots, we obtained curated expression data of 23819 individual captured locations on the ST array (Fig 1D). Unsupervised clustering and annotation using known liver cell markers were performed ^29,30^. The spatial spots were grouped into five major clusters: 1) Hepatocytes marked by *Alb, Cyp2e1,* and *Cyp1a2* expression; 2) Cholangiocytes marked by *Spp1, Epcam,* and *Krt19* (encode CK19) expression; 3) Erythrocytes express *Hba-a2, Hbb-bt,* and *Hba-bs*; 4) Stromal cells express *Col3a1, Col1a1* and *Col1a2*; and 5) Immune cells with *H2-Q6, H2-D1* and *Ccl5* expression (Figs 1D and S1D). It is noteworthy that a large proportion of spots (∼80%) from the ST array were associated with hepatocytes and cholangiocytes, while those cells only represent a small fraction (often less than 10%) for most marker-free liver scRNA-seq data (Fig S1E) ^5,31^. Hence, ST analysis is an ideal method for characterizing liver parenchymal cells.

The liver is constituted of highly structured repeating hexagonal anatomical units called liver lobules. Each liver lobule consists of six portal triads where bile ducts are located, a central vein, and space in between filled mainly with hepatocytes (Fig 1E). Hepatocytes in the liver are spatially and molecularly heterogeneous. They can be divided into pericentral, middle, and periportal zones organized along the center-to-edge axis, with hepatocytes performing dedicated metabolic functions in each zone – a phenomenon known as liver zonation (Fig 1E) ^32^. We further annotated hepatocyte zones using known zonation landmark genes ^30,33^. The annotated zones approximated hexagon shape, particularly in wild-type livers (Figs 1F and S1F-H). The zonation patterns of different livers had also been determined by staining E-cadherin (E-cad) and glutamine synthase (GS), markers of periportal and pericentral zones, respectively (Fig 1C). ST analysis and immunostaining results were largely consistent (Fig 1C, F). We observed that in *Nf2^−/−^;Wwc^+/–^*, *Nf2^−/−^;Sav1^+/–^*, and especially *Sav1^−/−^;Wwc^−/−^* livers, the pericentral zone was significantly shrunk, whereas the change in *Nf2^−/−^*, *Sav1^−/−^* or *WWC^−/−^* livers was not dramatic (Fig 1C, F, G). Moreover, the periportal zone in HPO1 inactivated livers (*Sav1^−/−^*, *WWC^−/−^*, and *Sav1^−/−^;Wwc^−/−^*) was expanded (Fig 1C, F, G). These results indicate that liver zonation is differentially perturbed following the inactivation of HPO1 and/or HPO2.

In ST analysis, a single spot may contain multiple cells. Recently, multiple algorithms have been developed to enhance the resolution of spatial transcriptome by coupling the scRNA-seq data ^34,35^. We perform deconvolution based on a public mouse liver single-cell dataset to increase spatial spot resolution and estimate cell-type compositions in each spot (GSE171993) ^5^. Shannon Entropy for each ST spot was calculated and generally low in hepatocyte zones. Hepatocyte zones had lower entropy, partly due to the large size of hepatocytes. However, Shannon Entropy for cells in portal areas, such as cholangiocytes and stromal cells, was relatively high, indicating the presence of different cells in these spots (Fig S1I). After decomposition, more cell types were identified, and these cells, mostly mesenchymal cells, localized mainly at portal areas. Cell types were changed dramatically in portal zones in mutant livers examined, whereas the hepatocyte zone of the complete HPO1 mutant liver (*Sav1^−/−^;Wwc^−/−^*) showed the most robust changes (Fig S1J, K). The heterogeneity and plasticity of hepatobiliary cells are further investigated in this study.

### Hippo signaling inactivation results in three pools of immature hepatobiliary cells

The change in abundance and location of hepatobiliary cells in livers with defective Hippo signaling suggests a perturbed liver development. To precisely characterize the states of hepatobiliary cells in different livers, we performed scRNA-seq for P21 *Nf2^−/−^;Wwc^+/–^* and *Sav1^−/−^;Wwc^−/−^* livers associated with drastic changes in ST analysis (Fig 1). Subsequently, we integrated our scRNA-seq data with a public single-cell atlas of developing mouse livers (Figs 2A and S2A-C). The P21 *Nf2^−/−^;Wwc^+/–^* and *Sav1^−/−^;Wwc^−/−^*livers contained three clusters of parenchymal cells which were different from normal P21 hepatobiliary cells: cluster 1 cells expressed high levels of embryonic hepatocyte markers *Afp, Igf2, H19,* and *Ahsg*; cluster 2 and 3 commonly expressed neonatal cholangiocyte markers *Peg3* and *Fos*. However, cluster 2 cells had high *Spp1, Lamc3, and Fxyd3 expression and intermediate Krt7 and Krt19 expression, whereas cluster 3 cells had high Krt7, Krt19,* and *Tspan8* expression and intermediate *Spp1* expression. Clusters 1, 2, and 3 cells exhibited immature features; for simplicity, they were termed as imHep, imCho1, and imCho2, respectively (Fig 2A, B).

**Figure 2.**
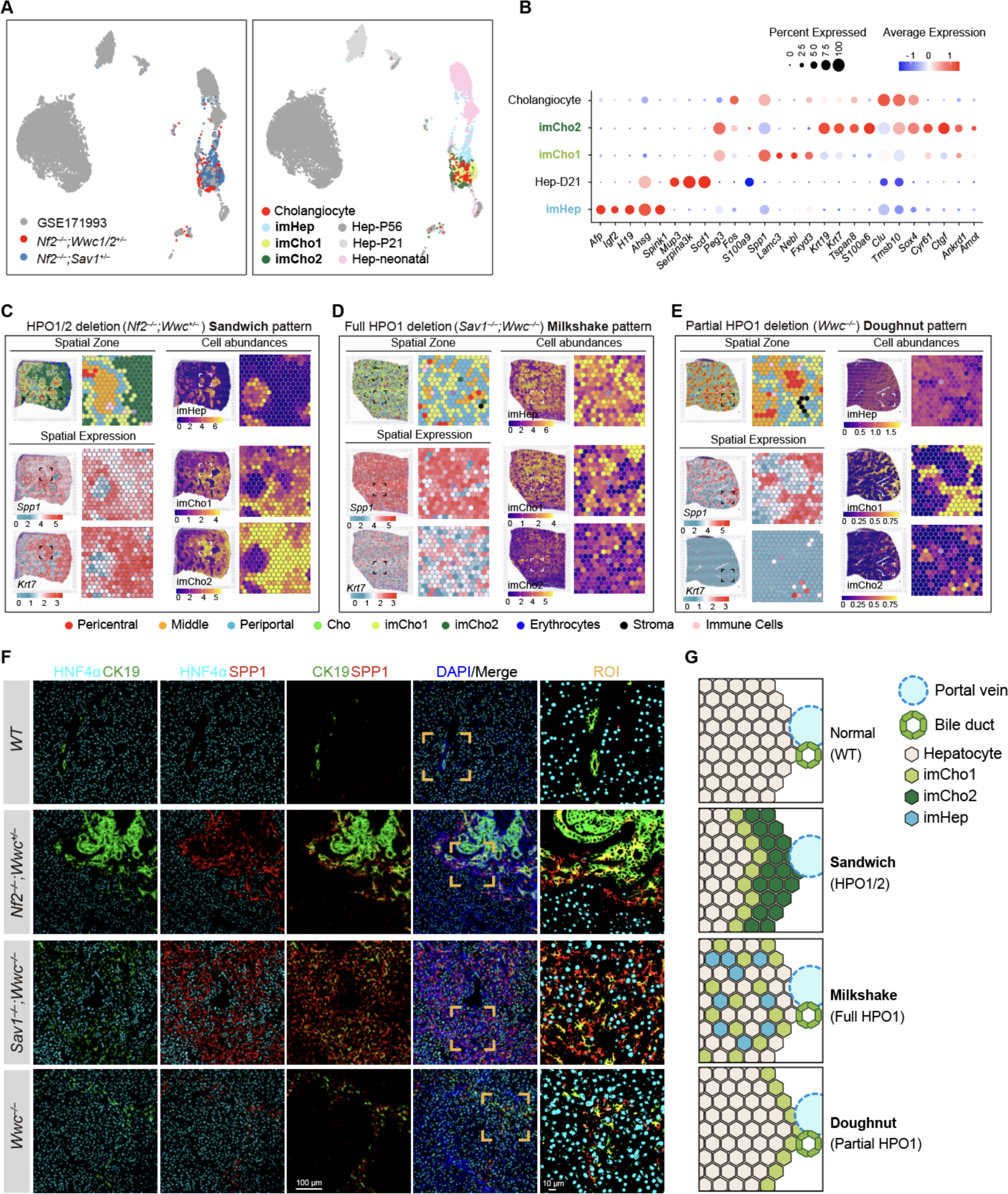
Three pools of immature hepatobiliary cells are associated with defective Hippo signaling. (A) UMAP representation of integrated hepatobiliary cells. The integrated cell atlas was generated by combining public scRNA-seq (GSE171993) and scRNA-seq data of *Nf2^−/−^;Wwc^+/–^* (HPO1/2) and *Wwc^−/−^;Sav1^-/–^* (full HPO1) livers. (B) Expression of selected markers for cell types. Dot size and color indicate the proportion of cells expressing marker genes in each cluster and the average gene expression level, respectively. (C-E) Distinct spatial distributions of imHep, imCho1, and imCho2 cells and their association with HPO1 and/or HPO2. The spatial organization of immature hepatobiliary cells is categorized into three types: Doughnut, Milkshake, and Sandwich. The expression of immature hepatobiliary cell markers and the abundance of each cell type at specific locations are projected. (F) Location of imHep, imCho1, and imCho2 cells. Tissue sections from livers with different genotypes are stained for HNF4α, SPP1, and CK19. Scale bar: 100 μm. (G) Diagram illustrating the spatial patterns of imHep, imCho1, and imCho2 cells across various livers with defective Hippo signaling.

We then analyzed the quantity and distribution of imCho1, imCho2, and imHep cells in livers with defective Hippo signaling. It appeared that imCho1 cells were commonly found in all livers with defective Hippo signaling, imCho2 cells were mainly present in livers with HPO2 inactivation (*Nf2^−/−^, Nf2^−/−^;Wwc^+/–^*, and *Nf2^−/−^;Sav1^−/−^*), and imHep cells were most robustly upregulated in livers with complete inactivation of HPO1 (*Sav1^−/−^;Wwc^−/−^*) livers (Figs 2C-E and S2D, E). Moreover, the spatial distribution of imCho1, imCho2, and imHep cells was highly patterned. In livers with both HPO1 and HPO2 inactivation, parenchymal cells were separated into three layers, with imCho1 cells residing between hepatocytes and imCho2 cells, and this pattern was designated as Sandwich (Figs 2C and S2D, E). Similar findings were observed in *Nf2^−/−^* livers, especially in areas with bile duct hamartomas (Fig S2D, E). In livers with complete HPO1 inactivation (*Sav1^−/−^;Wwc^−/−^*), imCho1 and imHep cells were evenly distributed and blended throughout liver parenchyma, and this pattern was designated as Milkshake (Figs 2D and S2D, E). In livers with partial HPO1 inactivation (*Sav1^−/−^* or *Wwc^−/−^*) or HPO2 inactivation (*Nf2^−/−^*), imCho1 cells were located along the edges of the hexagonal lobule and in close contact with periportal hepatocytes, and the pattern was designated as Doughnut (Figs 2E and S2D, E). Furthermore, the spatial and topological patterns of imCho1and imCho2 cells were also confirmed by immunostaining of CK19 and SPP1 (CK19^low^ SPP1^high^ and CK19^high^ SPP1^low^ indicate imCho1 and imCho2, respectively) (Figs 2F and S2F). Together, these findings suggest that imCho1, imCho2, and imHep cells are localized at distinct regions, have unique topological features, and are associated with activities of HPO1 and/or HPO2 (Fig 2G). During mouse liver development, cell migration mainly occurs during the early embryonic stage; as the liver gradually matures, the positions of hepatoblasts become relatively static, and differentiated hepatocytes and cholangiocytes are spatially separated. Hence, the featured distribution and topological patterns suggest that imCho1, imCho2, and imHep cells are likely derived from different progenitors.

### Inactivation of HPO1 blocks hepatocyte maturation and induces imHep cells

Hepatoblasts are present in both embryonic and neonatal livers and are marked by the expression of genes such as *Afp, Igf2, H19,* and *Ahsg* ^5^. These hepatoblast markers were highly expressed in imHep cells in *Sav1^−/−^;Wwc^−/−^* livers (Figs 2B, 3A and S3A). In addition, the expression of *Spink1*, another immature hepatocyte marker, was substantially increased in imHep cells and some hepatocytes in livers with partial HPO1 inactivation (Fig S3B) ^36^. The presence of AFP+ cells in *Sav1^−/−^;Wwc^−/−^* liver was also confirmed by immunostaining (Fig 3B). Hence, imHep cells have features similar to immature hepatocytes.

**Figure 3.**
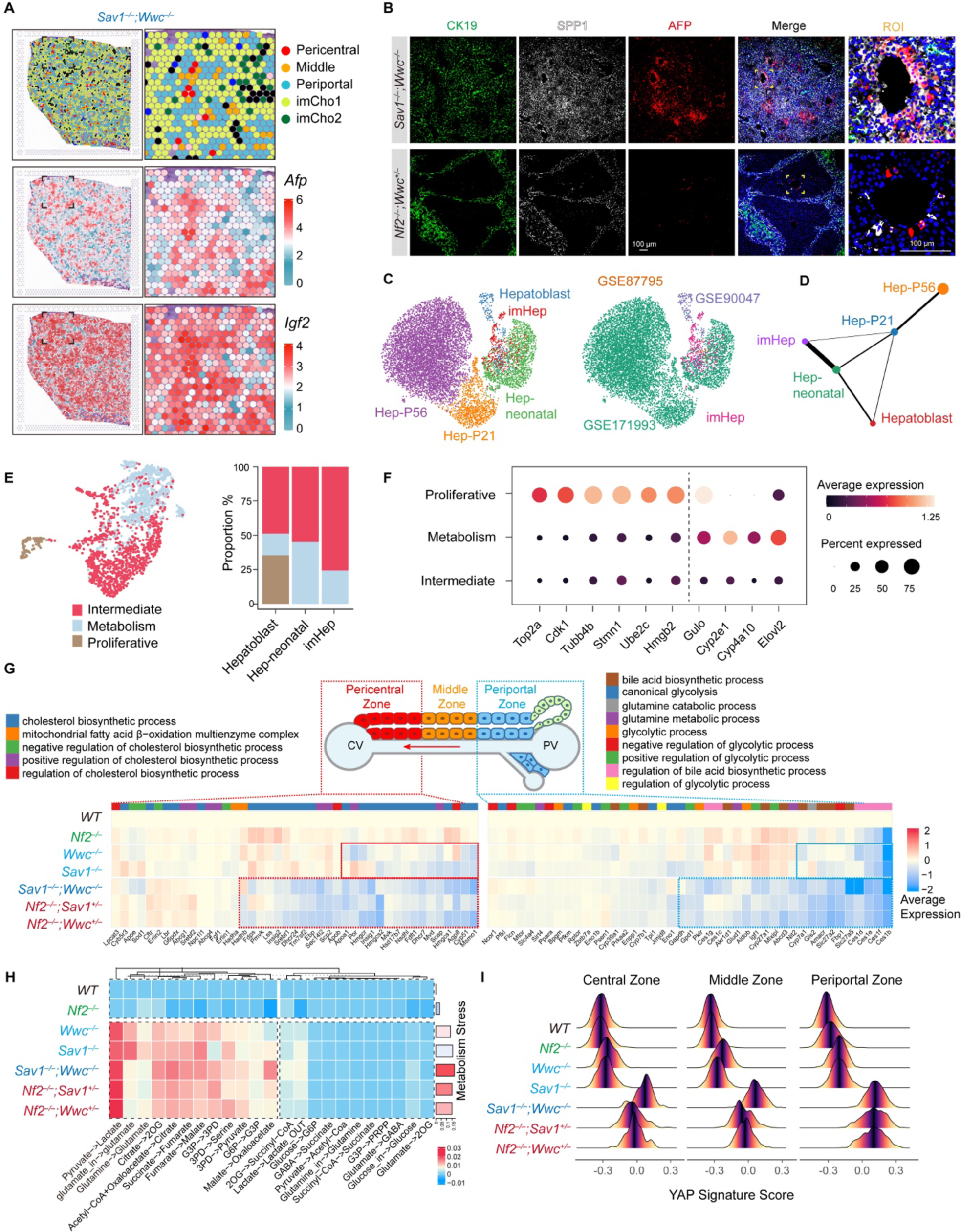
HPO1 regulates the postnatal maturation of hepatocytes. (A) The spatial expression of hepatoblast markers (*Afp* and *Igf2*) in *Wwc^−/−^;Sav1^−/−^* liver. (B) Immunostaining of AFP, SPP1 and CK19 in *Wwc^−/−^;Sav1^−/−^* and *Nf2^−/−^;Wwc^+/–^* livers. Scale bar: 100 μm. (C) UMAP representation of integrated scRNA-seq data. Left: cells are colored by cell type. Right: cells are colored by batch. Neonatal hepatocytes (P1, P3, P7) and embryonic hepatoblasts are clustered closely with imHep cells. (D) PAGA graph of different hepatocytes. The weight of the edges indicates statistical confidence between cell groups (nodes). (E) Sub-clustering analysis of immature hepatocytes. Immature hepatocytes can be divided into 3 clusters: metabolism, proliferative, and intermediate, according to the expression of their markers. The proportion of each cluster in different cell types is shown on the right. (F) Dot plot showing marker gene expression for clusters in (E). (G) Expression of metabolic genes in pericentral and periportal hepatocytes. A heatmap is constructed using spatial transcriptomics data of livers with defective Hippo signaling. (H) Heatmap of flux deviation normalized to the WT liver. A high value indicates a disbalance of cellular metabolism. The bar plot on the right shows the overall flux deviation of different livers. (I) Ridge plot of YAP signature score in hepatocyte zone. The x-axis represents the calculated YAP signature score obtained from hepatocyte zones.

To determine the developmental stage associated with imHep cells, we mapped imHep cells to hepatocytes atlas of developing (fetal to P56) mouse livers (GSE171993, GSE87795, GSE90047) ^5,31,37^. Integrated UMAP embedding showed that imHep cells deviated from P21 and P56 hepatocytes but clustered with neonatal and embryonic hepatoblasts (Fig 3C). Moreover, as indicated in partition-based graph abstraction (PAGA) analysis ^38^, imHeps cells were most closely related to neonatal hepatocytes (P1 to P7) (Fig 3D). Furthermore, three groups of hepatocytes (proliferative, metabolism, and intermediate) were identified as immature hepatocytes (Fig 3E, F). Intermediate and metabolism groups were commonly present in hepatoblasts, imHep cells, and neonatal hepatocytes, whereas the proliferative group was presented mainly in embryonic hepatoblasts (Fig 3E). These results suggest that the accumulation of imHep cells in full HPO1 inactivated livers was likely due to the retention and expansion of neonatal hepatocytes.

The liver functions as a central metabolism hub in the body. The metabolic function and zonation of hepatocytes are gradually established postnatally ^4,5^. The expression of hepatoblast markers and zonation defects suggest that the maturation of hepatocytes with HPO1 inactivation was delayed, and these livers may not effectively perform essential metabolic functions. We then estimated the expression of genes involved in metabolic functions, including bile acid biosynthesis, glycolytic process, and glutamine metabolism in the pericentral zone, as well as the cholesterol biosynthesis and β-oxidation function in the periportal zone ^32,39^. Indeed, compared to HPO2 inactivation, HPO1 and HPO1/2 inactivated livers exhibited significantly reduced expression of metabolic signature genes in both zones (Fig 3G). We also calculated single-cell metabolic flux in different livers ^40^. The results indicated that, except for *Nf2^−/−^*, livers with defective Hippo signaling were associated with high metabolic stress, indicating inefficient metabolic flux (Fig 3H). Together, these findings demonstrate the robust role of HPO1 in regulating the functional maturation of hepatocytes in postnatal development.

Next, we assessed whether YAP/TAZ activity was differentially regulated in hepatocytes by HPO1 and HPO2. YAP/TAZ activity in each hepatocyte zone was evaluated using a YAP/TAZ signature ^41^. Indeed, complete HPO1 deficiency (*Sav1^−/−^;Wwc^−/−^*) induced YAP/TAZ hyperactivation in all hepatocyte zones. HPO1/2 and partial HPO1 deficiency resulted in high and intermediate YAP/TAZ activation, respectively, with a dramatic effect observed in the periportal zones. On the other hand, HPO2 inactivation in hepatocytes had the weakest effect on YAP/TAZ activity (Figs 3I and S3C). Hence, the accumulation of imHep cells in livers with defective Hippo signaling is associated with YAP/TAZ activation.

### Inactivation of HPO1 and/or HPO2 promotes transdifferentiation of hepatocytes into imCho1 cells

The spatial location of imCho1 cells was predominant at the periphery of liver lobules or sandwiched between hepatocytes and imCho2 cells (Fig 2C-G). Hence, imCho1 cells might have originated from periportal hepatocytes or cholangiocytes. To infer the evolution and origin of the imCho1 population, we performed RNA velocity analysis, which yielded high-dimension vectors indicating the path and direction of cell state transition ^42,43^. The calculated flow vectors suggested that imCho1 can be derived from the imCho2 or imHep population, with more continuous flow vectors observed from imHep to imCho1 (Fig 4A). Indeed, imCho1 expressed markers of cholangiocytes derived from hepatocytes, such as *Spp1*, *Ccl2*, *Vim*, and *Ncam1* (Fig 4B) ^44-46^. Consistent with bioinformatic analysis, we observed cells retained a classical hepatocyte nuclei morphology yet exhibited a decreased HNF4α expression and elevated SPP1 expression in *Sav1^−/−^;Wwc^−/−^* livers (Fig 4C, #2). These cells likely represented a transition state from hepatocytes to imCho1. To validate if imCho1 cells were derived from a hepatocyte origin, we deleted HPO1 and/or HPO2 genes directly in hepatocytes. Adeno-associated virus serotype 8 expressing Cre recombinases under the hepatocyte-specific thyroxine-binding globulin (*Tbg*) promoter (AAV8-TBG-Cre) was administrated to P28 wild type (WT), *Sav1^F/F^;Wwc^F/F^*, or *Nf2^F/F^;Wwc^F/F^* Ai9 mice and livers were harvested for examination at indicated time points (Figs 4D and S4A, B). In these mice, tdTomato would be expressed following Cre recombination and used for lineage tracing. Livers were normal at seven days following injection; however, after 14 days, *Nf2^F/F^;Wwc^F/F^* and *Sav1^F/F^;Wwc^F/F^*livers showed robust enlargement, bile duct expansion, AFP expression, and disrupted zonation (Fig S4A-D). In these livers, we observed normal hepatocytes (#0: SPP1– and CK19–; tdTomato-positive) and normal cholangiocytes (#1: SPP1+ and CK19+; tdTomato-negative). Meanwhile, we also observed tdTomato-positive hepatocytes (#2, SPP1+ and CK19–) and cholangiocytes (#3, SPP1+ and CK19+); these hepatocytes-derived cells likely represented imHep and imCho1 (Fig 4E). Hence, imHep and imCho1 cells may be derived from hepatocytes via dedifferentiation and transdifferentiation, respectively. Hepatocytes may be directly transdifferentiated into imCho1 cells or dedifferentiated into imHep cells and reprogrammed into imCho1 cells (Fig 4F). YAP has been shown to induce the expression of transcription factor BCL3, and the latter is critical in mediating hepatocytes to cholangiocytes transition ^47^. Indeed, the expression of *Bcl3* was highest in imCho1 cells (Fig S4E), which might be involved in the formation of imCho1 cells.

**Figure 4.**
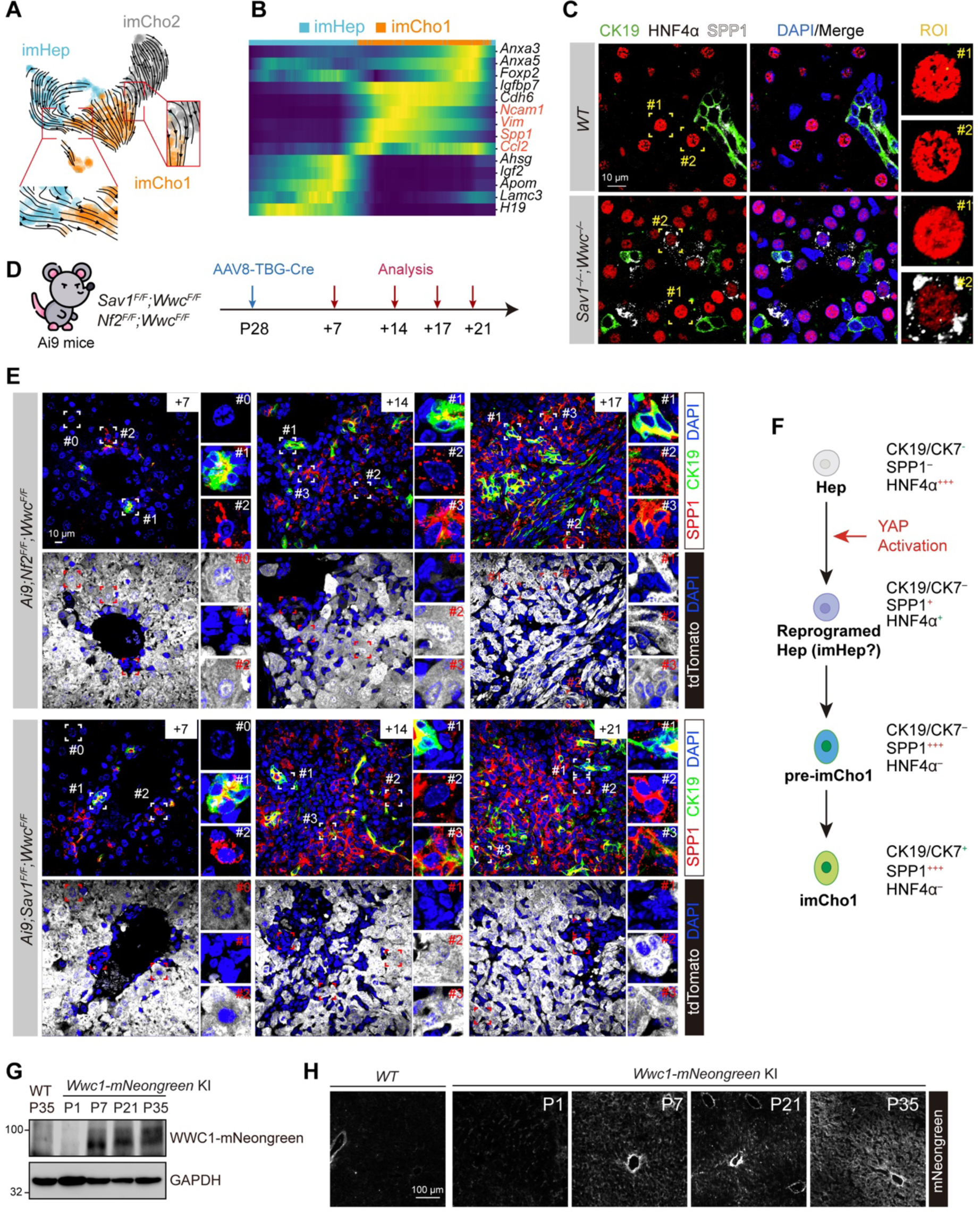
imCho1 is derived from hepatocyte transdifferentiation. (A) RNA velocity streamline plot depicting potential cell transitions. Streamline number and continuity reflect the predicted probability of cell fate transitions. (B) Heatmap of gene expression during hepatocyte transdifferentiation into imCho1. (C) Immunostaining of CK19, HNF4α, and SPP1 expression in liver sections. 3-week-old control or *Sav1^−/−^;Wwc^−/−^*livers are analyzed. WT livers display high HNF4α expression and no SPP1 expression in all hepatocytes (ROI #1 and ROI #2). In *Sav1^−/−^;Wwc^−/−^* livers, in addition to normal hepatocytes (ROI #1), a population of hepatocytes with low HNF4α and high SPP1 expression (ROI #2) emerges. Scale: 10 μm. ROI: region of interest. (D) Schematic diagram showing hepatocyte-specific Hippo pathway gene deletions and lineage tracing experiment. Low-dose AAV8-TBG-Cre is injected via tail vein into 4-week-old mice with genetic backgrounds of *Ai9;Nf2^F/F^;Wwc^F/F^*and *Ai9;Sav1^F/F^;Wwc^F/F^*. Liver samples were harvested and analyzed at 7 days (+7), 14 days (+14), 17 days (+17), or 21 days (+21) after injection. Hepatocytes with Cre expression are traced by tdTomato expression. (E) Immunostaining of SPP1, CK19, and tdTomato in liver sections. Livers from mice at different time points following AAV8-TBG-Cre injection are collected, sectioned, and subjected to immunostaining. For each genotype, the upper panels are merged signals from SPP1 (red), CK19 (green), and DAPI (blue); lower panels show tdTomato (white) and DAPI (blue) for the same field. ROI #0: Normal hepatocytes (no SPP1 expression, tdTomato positive); ROI #1: Cholangiocytes and cholangiocyte-derived cells (low SPP1 expression, high CK19 expression, tdTomato negative); ROI #2: Pre-imCho1 cells (SPP1 positive, tdTomato positive); ROI #3: imCho1 cells derived from hepatocytes (high SPP1 expression, low CK19 expression, tdTomato positive). Scale bar: 10 μm. (F) Schematic diagram showing hepatocyte reprogramming and stepwise transdifferentiation into imCho1. (G and H) Temporal analysis of the expression of *Wwc1*-mNeongreen. Liver samples were obtained from WT mice and *Wwc1*-mNeongreen knock-in mice on different postnatal days (1, 7, 21,35). Protein expression (G) and immunostaining with TSA (H) are shown.

When comparing the phenotypes of hepatocytes-specific knockouts (AAV8-TBG-Cre), we noticed more imHep and imCho1 cells in *Sav1^F/F^;Wwc^F/F^*livers than that in *Nf2^F/F^;Wwc^F/F^* livers (Figs 4E and S4C). However, more bile ducts and robustly enlarged gallbladders were observed in *Nf2^F/F^;Wwc^F/F^*livers, and these mice were sick and euthanized 17 days after injection (Fig S4A, B). These differences suggest that SAV1 (HPO1) and NF2 (HPO2) play dominant roles in hepatocytes and cholangiocytes, respectively. The expression of WWC1 and SAV1, two HPO1 components, are kept at low levels in early postnatal mouse livers and induced after weaning ^25^. To monitor cell type-specific expression of WWC1, we engineered conditional knockin mice, which, when crossed with *Alb*-cre mice, express WWC1-mNeonGreen fusion protein, specifically in hepatobiliary cells (Fig S4F). Interestingly, the expression of WWC1-mNeonGreen was absent in P1 livers, appeared in P7 livers, and gradually elevated in P21 and P35 livers, as indicated by both immunoblotting and immunostaining (Fig 4G, H). In a detailed examination of full HPO1 inactivated livers (*Sav1^−/−^;Wwc^−/−^*), the expansion of imCho1 and imHep cells were not evident in early postnatal livers (P1-P7) but robustly induced in P21 livers (Fig S4G). Hence, the HPO1 signaling in mouse liver is established gradually when hepatocytes are undergoing functional maturation, and inactivating HPO1 leads to retention of neonatal hepatocytes (imHep) and transdifferentiation to immature cholangiocytes (imCho1). HPO1 likely serves as a postnatal maturation checkpoint in hepatocytes to suppress YAP/TAZ activity, cell plasticity, and organ growth.

### Inactivation of HPO2 blocks cholangiocyte maturation and induces imCho2 cells

During embryonic liver development, hepatoblasts in contact with the portal vein form a ductal plate structure containing two layers of immature cholangiocytes from E14-E18 (Fig 5A) ^48^. In the perinatal stage, the ductal plate undergoes a remodeling process involving apoptosis and tubulogenesis to form intrahepatic bile ducts ^49^. We integrated single-cell data of mouse livers (E9.5-E17.5) containing hepatoblasts, cholangiocytes, erythrocytes, endothelial cells, macrophages, and megakaryocytes and located these cells spatially by performing deconvolution against a spatial transcriptome of E16.5 mouse livers (Fig 5B) ^50^. Interestingly, E16.5 cholangiocytes were clustered and exhibited structures similar to ductal plates and expressed imCho2 markers (Fig 5B, C). Furthermore, SCENIC analysis also indicated similarities between E16.5 cholangiocytes with imCho2 cells (Figs 5C and S5A).

**Figure 5.**
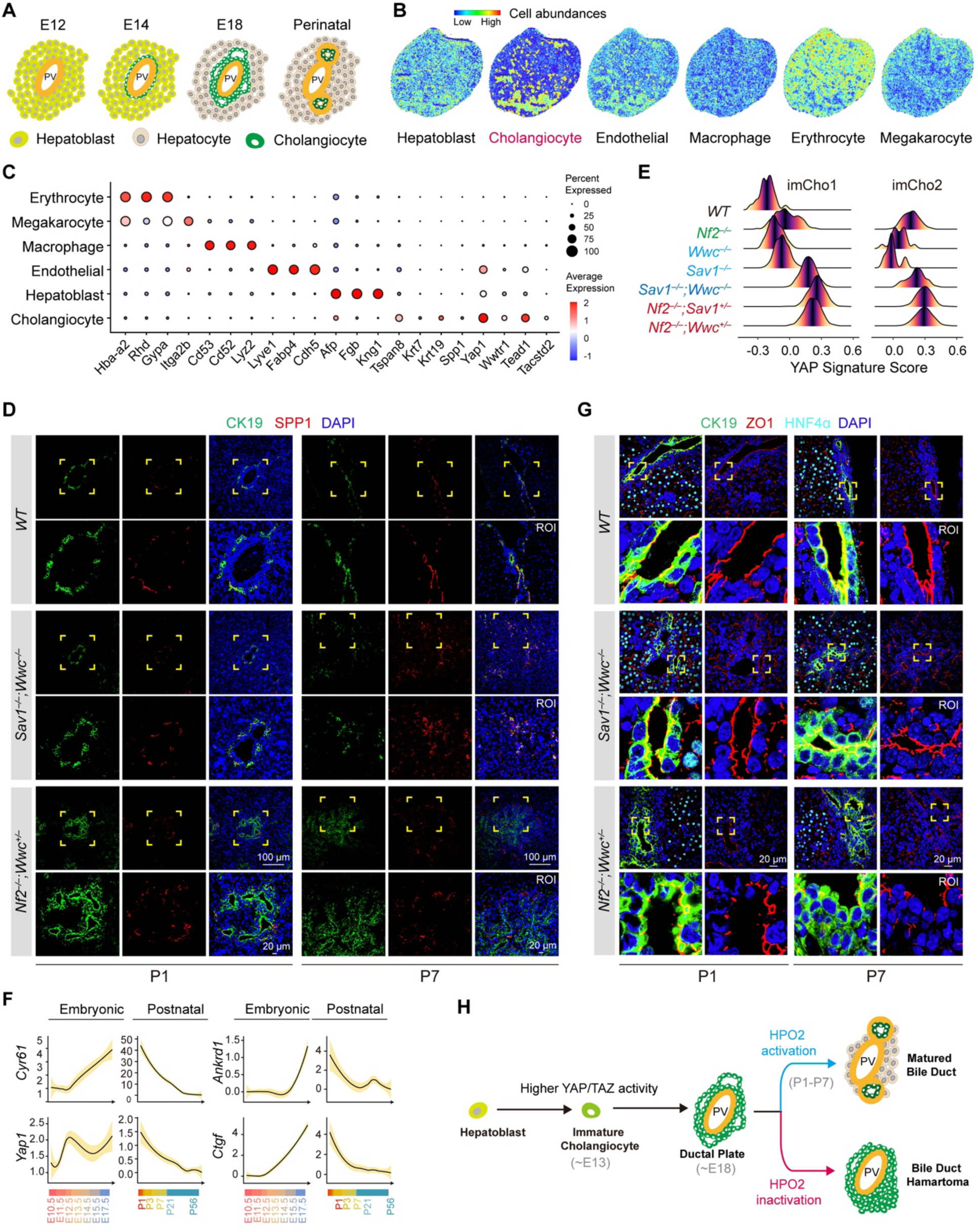
HPO2 mediates ductal plate remodeling and maturation of cholangiocytes. (A) Embryonic and perinatal development of bile ducts in mice. Initially, hepatoblasts surrounding the portal vein (PV) commit into cholangiocyte lineage and form a single-layered ring structure by embryonic day 14 (E14). These immature cholangiocytes proliferate to form a bilayer structure called the ductal plate at E18. At birth, this ductal plate undergoes remodeling and regression to form mature and functional bile ducts (B) Spatial distribution of different cell types in E16.5 mouse liver. The cell abundances are calculated by deconvolution analysis using stereo-seq data and scRNA-seq data from mouse fetal livers. (C) Expression of selected genes in cells at the ductal plate stage. E16.5 cholangiocytes express high *Yap*, *Taz* (*Wwtr1*), *Tspan8*, and *Krt19*. Conversely, *Spp1* expression is low. The expression pattern is similar to that of imCho2 cells. (D) Immunostaining of CK19 and SPP1 in different liver sections. Livers from WT, *Sav1^−/−^;Wwc^−/−^*, and *Nf2^−/−^;Wwc^+/–^* mice at P1 and P7 are analyzed. Yellow boxes indicate ROI. Scale: 100 μm and 20 μm. (E) Ridgeline plot showing YAP/TAZ activity in cholangiocytes from mice with different genotypes. The horizontal axis represents relative YAP/TAZ activity. (F) Integration and analysis of scRNA-seq data of cholangiocytes at different developmental stages. During the perinatal period, the expression of *Yap* and its target genes gradually increased and peaked at embryonic day 17.5 (E17.5). After birth, as the bile duct plate continuously regresses, the expression of *Yap* and its target genes gradually decreases. (G) Immunostaining of ZO1, CK19, and HNF4α in different liver sections. The same samples are used in D. Yellow boxes indicate ROI. Scale: 20 μm. (H) Diagram illustrating how HPO2 regulates the commitment and maturation of cholangiocytes by regulating YAP/TAZ activity. Inactivation of HPO2 leads to failed ductal plate remodeling and the formation of bile duct hamartomas.

We then assessed the ductal remodeling in neonatal mice with HPO1 and/or HPO2 inactivation. Ductal plate structures were evident in all P1 livers analyzed, as indicated by CK7 and CK19 positive cells arranged in two layers (Figs 5D and S5B). At P7, ductal plate structures were regressed in WT and *Sav1^−/−^;Wwc^−/−^*livers, but further dilated in *Nf2^−/−^;Wwc^+/–^*livers (Figs 5D and S5B). In P1 *Nf2^−/−^;Wwc^+/–^* livers, most CK19+ cells were surrounded by SPP1+ cells, and these cells might be imCho2 and imCho1 cells respectively (Fig 5D). The juxtaposition of these cells was identical to the Sandwich pattern shown above (Fig 2E-I). Hence, the imCho2 cells are likely perinatal ductal cells that have failed to undergo ductal remodeling; the proliferation and accumulation of these cells result in the dilation or expansion of ductal plates in portal areas.

Besides *Nf2^−/−^;Wwc^+/–^* and *Nf2^−/−^;Sav1^+/–^*livers, the accumulation of imCho2 cells was also present in hamartomas in *Nf2^−/−^*livers (Fig 2A,C). Hence, imCho2 cells appear to be associated with HPO2 inactivation. We used ST data to compare the YAP/TAZ signature in imCho2 cells and observed strong YAP/TAZ activation in full HPO1 and HPO1/2 perturbed livers. YAP/TAZ activation was also significantly induced in *Nf2^−/−^*cells. The YAP/TAZ signature change in imCho1 cells followed a similar trend (Figs 5E and S5C). Hence, YAP/TAZ activity in cholangiocytes and periportal hepatocytes might be more sensitive to HPO2 inactivation, which contributes to the accumulation of immature cholangiocytes, especially imCho2 cells, in HPO2 defective livers.

We also engineered conditional knockin mice to express NF2-mNeonGreen fusion protein in hepatobiliary cells (Fig S5D). In contrast to the age-dependent expression of WWC1, the expression of NF2 in hepatobiliary cells was constant at different postnatal stages (Fig S5E, F). It has been shown previously that YAP/TAZ is required to form ductal plates, especially the second layer of cholangiocytes ^51^. By integrating and analyzing scRNA-seq data of cholangiocytes from embryonic stage to maturation (GSE87795 and GSE90047), we discovered that YAP/TAZ activity in cholangiocytes, as indicated by the expression of *Cyr61*, *Ctgf*, and *Ankrd1*, was gradually increased during embryonic development (Fig 5F). However, the expression of these genes immediately declined after birth (Fig 5F). Hence, the Hippo signalings, especially HPO2, may be up-regulated to promote ductal plate regression by restricting YAP/TAZ activity. NF2 is a component of cell-cell junctions, and its junctional localization is required for regulating Hippo signaling ^52^. Ductal plate regression and bile duct tubulogenesis may form strong cell-cell junctions, which promote the junctional localization of NF2 and activation of HPO2, leading to bile duct maturation. In wild-type livers, tight junctions of P7 cholangiocytes, as indicated by ZO-1 staining, were strong and more continuous than those of P1 cholangiocytes. Interstingly, tight junctions were disrupted in P1 and P7 cholangiocytes from *Nf2^−/−^;Wwc^+/–^* livers. On the other hand, the change in tight junction was less dramatic in *Sav1^−/−^;Wwc^−/−^*cholangiocytes (Fig 5G). It is likely that, during perinatal liver development, HPO2 serves as a checkpoint for YAP/TAZ activity in cholangiocytes, and reduction in YAP/TAZ activity is required for ductal plate regression and bile duct maturation. In HPO2-inactivated livers, cholangiocytes will retain features of the ductular plate and keep proliferating, leading to the accumulation of immature cholangiocytes (imCho2) in portal areas and the formation of bile duct hamartomas or cholangiocarcinomas (Fig 5H).

### *Yap/Taz* knockout accelerates liver maturation and damage

Ectopic *Yap* expression or YAP/TAZ activation leads to liver enlargement (Fig 1) ^14,18^. However, as documented by several studies, *Yap^−/−^;Taz^−/−^* livers are nearly normal or slightly larger than WT livers ^21-24^. It is currently a challenge to address these counterintuitive phenotypes resulting from the gain or loss of YAP/TAZ functions, and the role of YAP/TAZ in cell proliferation and organ size control has been questioned ^22,23^. Since inactivations of Hippo signaling in livers led to a robust accumulation of immature cells and delayed liver functional maturation (Figs 2-5), we asked whether *Yap/Taz* deletion affects the maturation of hepatobiliary cells.

To investigate the role of YAP/TAZ in perinatal and postnatal liver development, we harvested WT and *Yap^−/−^;Taz^−/−^* (*Alb*-cre) livers at various stages (P1, P7, P12, P21, and P28). The gross appearance and liver/body weight ratios of *Yap^−/−^;Taz^−/−^*livers were similar to that of WT livers (Fig S6A, B). Consistent with previous studies, cholangiocytes (CK19+) and ductal progenitors (SOX9+) were present at portal areas in WT livers, whereas they dramatically reduced in *Yap^−/−^;Taz^−/−^* livers (Fig S6C, D) ^21-24^. During postnatal development, livers gradually turn off liver hematopoiesis and the expression of fetal genes and switch on the adult gene program to carry out metabolic functions ^53^. Strikingly, the number of erythroblasts, an indicator of liver hematopoiesis, was significantly reduced in P7 and P12 *Yap^−/−^;Taz^−/−^* livers (Fig 6A, B). The expression of fetal genes, such as *Afp*, was dramatically reduced in P7 and P12 *Yap^−/−^;Taz^−/−^* hepatocytes (Fig 6C, D). Moreover, significantly more binucleated hepatocytes were detected in P1 and P7 *Yap^−/−^;Taz^−/−^* livers (Fig S6E, F). Furthermore, GS+ hepatocyte clusters were detected in P1 and P7 *Yap^−/−^;Taz^−/−^*livers, indicating an early establishment of metabolic functions (Fig 6E, F). These data suggest early hepatocyte maturation in *Yap^−/−^;Taz^−/−^* livers.

**Figure 6.**
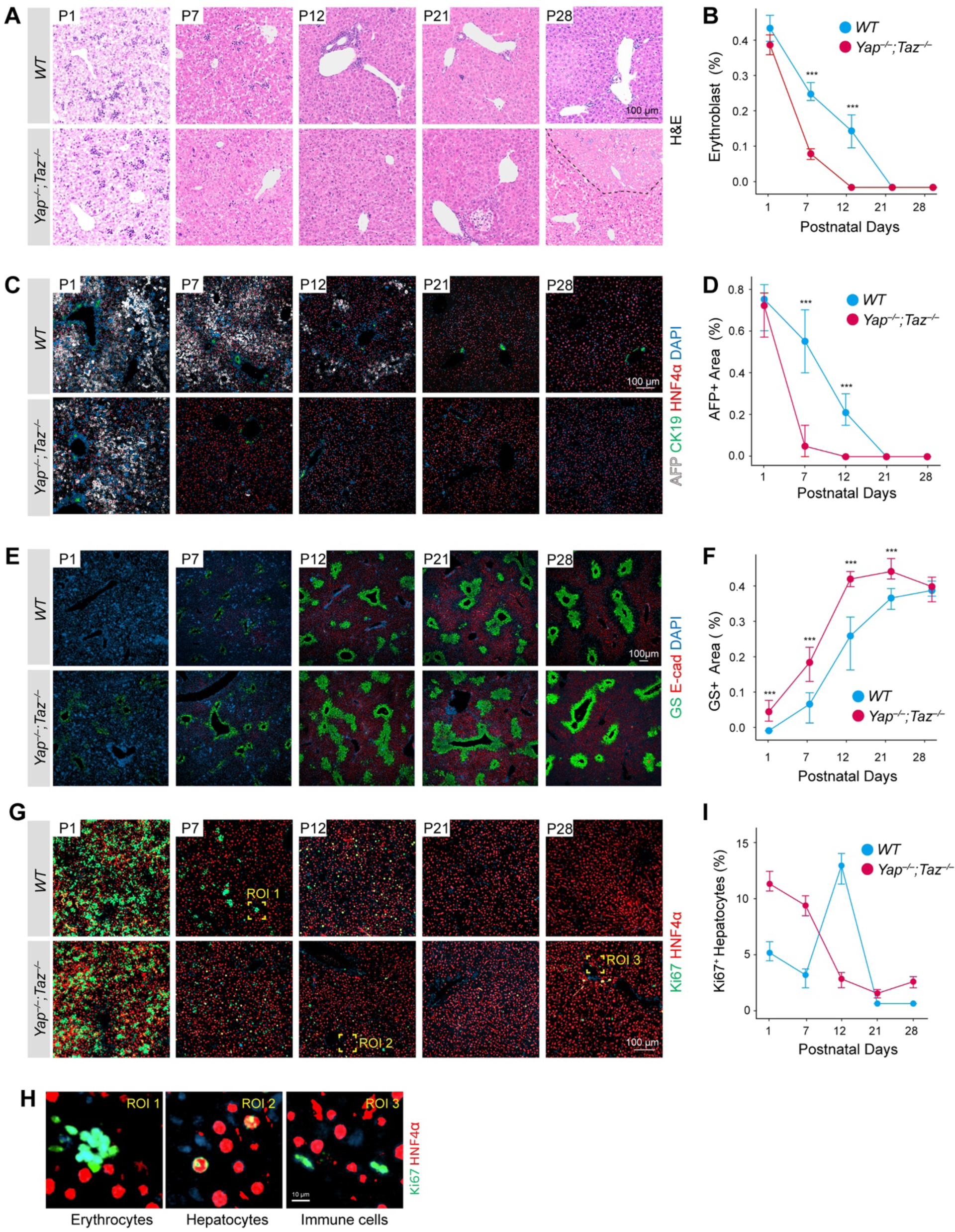
*Yap/Taz* deletion results in precocious liver maturation and dysfunction. (A and B) Liver-specific *Yap/Taz* deletion induces cessation of liver hematopoiesis and hepatic necrosis. H&E staining of liver sections from WT and *Yap^−/−^;Taz^−/−^* mice at postnatal days (P) 0, 7, 12, 21, and 28. Compared to WT livers, *Yap/Taz* deficient livers showed significantly fewer erythroblasts at P7 and evident necrotic areas at P28. Scale bar: 100 μm. Statistical analysis is shown in (B). (C and D) *Yap/Taz* deletion reduces AFP expression in neonatal livers. Immunostaining of AFP, CK19, and HNF4α in liver sections from WT and *Yap^−/−^;Taz^−/−^*mice. Hepatocytes in *Yap/Taz* deficient mice display a marked decrease in AFP expression at P7 and P12. Scale bar: 100 μm. Statistical analysis is shown in D. (E and F) *Yap/Taz* deficient livers exhibit abnormal zonation. Immunostaining of E-cadherin (Ecad) and glutamine synthetase (GS) in WT and *Yap^−/−^;Taz^−/−^* livers. *Yap/Taz* deficient hepatocytes exhibit markedly increased GS expression at P7 and P12. Scale bar: 100 μm. Statistical analysis is shown in (E). (G-I) *Yap/Taz* deletion disturbs hepatocyte proliferation. Immunostaining of Ki67 and HNF4α in WT or *Yap/Taz* deficient mouse liver sections. Proliferative hepatocytes (Ki67 and HNF4α double positive) in *Yap/Taz* deficient livers are more at P1 to P7 and less at P12. Scale bar: 100 μm. High-magnification views of ROI are shown in H. Scale bar: 10 μm. Statistical analysis is shown in I. Data are presented as mean ± SEM from at least three independent biological replicates.

The Hippo pathway has been thought to regulate both hepatocyte proliferation and cell death, which determine cell numbers and liver size ^54^. The proliferation of hepatocytes during liver development is highly dynamic, with P12 hepatocytes having the highest proliferation rate ^4^. We observed similar results in the WT liver, but the proliferation of *Yap^−/−^;Taz^−/−^* hepatocytes were high at P1 and P7 and relatively low at P12, indicating an ahead-of-schedule proliferation in *Yap^−/−^;Taz^−/−^* livers (Fig 6G-I). In P1 and P7 WT livers, most Ki67-positive cells are erythroblasts, and only Ki67 and HNF4α double-positive cells were counted as proliferating hepatocytes (Fig 6H). Moreover, necrotic areas were observed in P28 *Yap^−/−^;Taz^−/−^*livers, as indicated by H&E and TUNEL staining (Figs 6A and S6G, H). Liver damage may evoke a compensatory or regenerative response to induce the proliferation of hepatobiliary cells, and indeed, the proliferation of hepatocytes in P28 *Yap^−/−^;Taz^−/−^*livers were slightly higher than that of WT livers (Fig 6I). Thus, both proliferation and survival of hepatocytes are perturbed in *Yap^−/−^;Taz^−/−^* livers, even though the net effect on liver size is not evident.

Together, these data indicate a critical role of YAP/TAZ in regulating the fetal-to-adult transition in postnatal liver development. In the absence of YAP/TAZ, although the gross appearance of livers is nearly normal, the liver maturation program starts early and proceeds more rapidly. Moreover, the proliferation of hepatocytes was perturbed, and hepatocyte death was evident in the *Yap^−/−^;Taz^−/−^* livers (Fig 6). Hence, the Hippo signaling likely promotes postnatal liver maturation by suppressing YAP/TAZ activity, and YAP/TAZ deficiency promotes liver maturation and liver damage.

### Liver damages induce patterned YAP/TAZ activation and accumulation of immature cells

Unlike many other organs, the mammalian liver has a regenerative capacity following tissue damage ^8^. The Hippo pathway is also involved in liver regeneration ^23,46,55,56^. Immature cells associated with defective Hippo signaling arise following liver damage and participate in liver regeneration. We analyzed available datasets of regenerating livers following acetaminophen (APAP) or 3,5-diethoxycarbonyl-1,4-dihydrocollidine (DDC) treatments, which induce injuries of pericentral or periportal parenchymal cells, respectively^55,57^.

In livers treated with APAP, pericentral hepatocytes undergo rapid necrosis, and hepatocytes in remaining zones will proliferate and repopulate the liver gradually within 96 hrs; the spatiotemporal changes of livers in this process have been recently profiled using scRNA-seq and ST ^57^. In these datasets, a cluster of hepatocytes identical to imHep cells was identified, as indicated by high *Afp* and *Spp1* expression (Fig S7A, B). On the other hand, the presence of imCho1 and imCho2 cells was not evident (Fig S7A, B). Interestingly, a subset of hepatocytes expressing *Afp* at the regenerating front has been defined as interface hepatocytes ^57^. On the ST matrix, the expression of both YAP/TAZ signature genes and *Afp* was restricted to the boundaries of damaged areas (Fig S7A). In tissue sections from livers at different time points after APAP injection, AFP and SPP1 double-positive cells were identified at regenerating boarders in 24 hrs and 48 hrs livers but were absent in 0 hr (control) and 96 hrs (regenerated) livers (Fig S7C). A similar pattern was observed for YAP/TAZ expression (Fig S7D). Hence, upon pericentral liver damage, hepatocytes adjacent to damaged areas may upregulate YAP/TAZ activity to promote the transition of hepatocytes into an immature state similar to imHep cells.

DDC treatment induces injuries on both periportal hepatocytes and cholangiocytes, and the regenerating process features atypical ductal proliferation ^55^. As indicated by scRNA-seq analysis of EPCAM+ cells following DDC treatment, the cholangiocytes during regeneration were highly heterogeneous ^55,58^. We performed combinatory analysis on scRNA-seq datasets of cholangiocytes from livers treated with DDC (ChoDDC) or livers with perturbed Hippo signaling and observed similarities between ChoDDC and imCho2 (Fig S7E-G). We also stained tissue sections from regenerating livers treated with DDC and identified the expansion of CK19 and SPP1 expressing cells with imCho2 or imCho1 features at portal areas (Fig S7H). Together, these results suggest that, upon periportal injuries, cholangiocytes and periportal hepatocytes may be converted into immature states identical to imCho2 and imCho1, respectively, to facilitate liver regeneration.

Immature cells similar to imHep and imCho1/2 cells appeared in pericentral and periportal areas following APAP and DDC treatments, respectively (Fig S7). Since HPO1 and HPO2 play predominant roles in hepatocytes and cholangiocytes, they may be specifically perturbed by pericentral and periportal liver damage, and contribute to cell type-specific YAP/TAZ activation and tissue repair. It would be interesting to explore how HPO1 and HPO2 are dynamically regulated following liver damage.

## DISCUSSION

In this study, by combining genetic models and ST analysis, we have shown that HPO1 and HPO2 play predominant roles in hepatocytes and cholangiocytes, respectively. Inactivation of HPO1 and/or HPO2 leads to defective differentiation and maturation of hepatobiliary cells, resulting in robust expansion of immature cells. On the other hand, deletion of *Yap/Taz* promotes liver maturation. Hence, the Hippo signalings are spatiotemporally regulated during perinatal and postnatal liver development to supervise the maturation of hepatobiliary cells, and we propose that HPO1 and HPO2 serve as maturation checkpoints for hepatocytes and cholangiocytes, respectively, to ensure proper liver development and size control (Fig 7).

**Figure 7.**
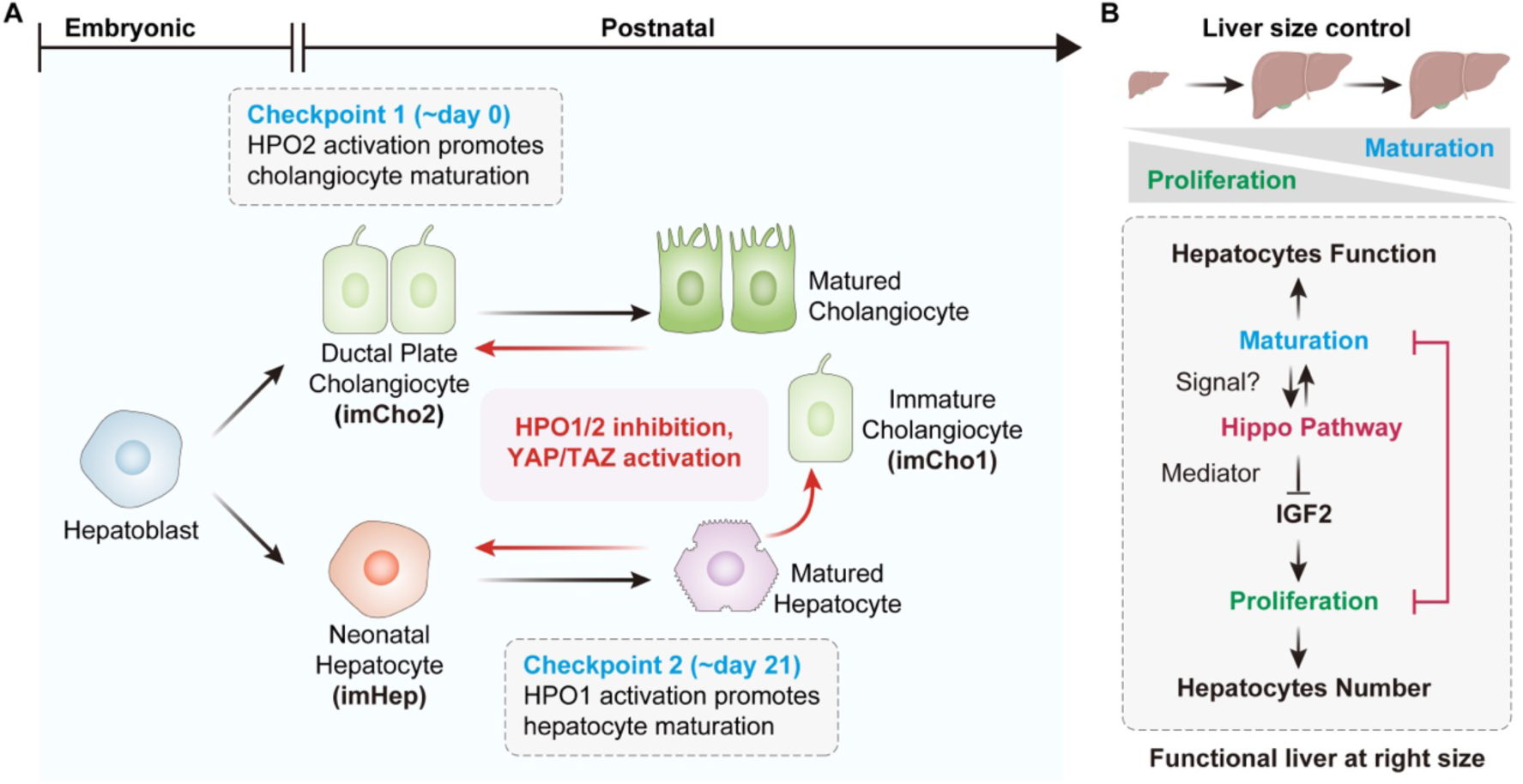
Spatiotemporal Hippo signalings instruct proliferation and maturation of liver parenchymal cells. (A) HPO1 and HPO2 function as maturation checkpoints for hepatocytes and cholangiocytes, respectively, during liver development. Higher YAP/TAZ activity promotes the specification of hepatoblasts into immature cholangiocytes during embryonic liver development. HPO1 is activated at 2-3 weeks postnatally to inhibit YAP/TAZ activity in hepatocytes and facilitate hepatocyte maturation. On the other hand, HPO2 is immediately activated after birth and is required for ductal plate remodeling and maturation of cholangiocytes. Inactivation of HPO1 leads to the accumulation of immature hepatocytes (imHep), transdifferentiation of hepatocytes into cholangiocytes (imCho1), and dramatic liver enlargement. Inactivation of HPO2 during liver development results in the accumulation of immature cholangiocytes (imCho2) and the formation of bile duct hamartomas. (B) Hepatocyte proliferation and maturation, both regulated by HPO1, determine hepatocyte number and function, ultimately leading to a fully functional liver at the right size. Hepatocyte maturation promotes the establishment of HPO1, which, in turn, further enhances cell maturation while inhibiting cell proliferation by suppressing IGF2 expression to restrict liver size. Understanding how HPO1 senses maturation presents an intriguing question for future study.

### Hippo signalings as liver maturation checkpoints

Hepatocytes are specified from bipotent hepatoblasts and gradually mature to gain metabolic functions in postnatal development ^5^. In livers with HPO1 or HPO2 inactivation, some hepatocytes, especially those at portal areas, are converted into immature cholangiocytes (imCho1), which is consistent with previous reports that activation of YAP drives hepatocytes to cholangiocytes transdifferentiation (Figs 2, 4) ^59^. Moreover, in the complete HPO1 inactivated (*Sav1^−/−^;Wwc^−/−^*) livers, accumulation of immature hepatocytes (imHep) is observed throughout the whole livers, and these cells are similar to neonatal hepatocytes (Fig 3). The phenotype of *Sav1^−/−^;Wwc^−/−^* liver is weak at P1 and P7, as indicated by sparse imCho1 and imHep cells, suggesting a low HPO1 activity in perinatal hepatocytes (Fig S4G). Indeed, the expression of WWC1, which is highly correlated to HPO1 activity, is low in perinatal hepatobiliary cells and gradually upregulated postnatally ^21,25^ (Fig 4). HPO1 appears to be specifically turned on when the liver size and maturation status reach a threshold, and HPO1 activation, in turn, limits YAP/TAZ activity and liver growth. Hence, HPO1, as a postnatal hepatocyte maturation checkpoint, may ensure proper live growth and functionality (Fig 7A).

In embryonic mouse liver development, a relatively higher YAP/TAZ expression is required for the specification of cholangiocytes from hepatoblasts and the formation of ductal plate structures ^51^. The high basal YAP/TAZ activity assigned to cholangiocytes may require additional regulation. Hence, HPO2 is designated to control YAP/TAZ activity in cholangiocytes. Perinatal ductal plate remodeling leads to the regression of ductal plate structures and tubulogenesis of bile ducts, which is essential for the maturation of cholangiocytes ^49^. HPO2 inactivation leads to a dramatic expansion of imCho2 cells similar to ductal plate cholangiocytes, indicating a critical role of HPO2 in cholangiocyte maturation. Ductal plate remodeling and bile duct maturation are accompanied by the formation of strong cell-cell junctions between cholangiocytes (Fig 5G). The formation of these junctions may serve as a signal to turn on HPO2 because the localization of NF2 to cell-cell junctions is required for its function in the Hippo pathway ^52,60^. Hence, HPO2 may be a perinatal cholangiocyte checkpoint to guide their maturation (Fig 7A).

The regulation of cell maturation by the Hippo pathway may also be present in additional organs. Recently, it has been shown that YAP activity is high in the fetal intestine and low in the adult intestine, and YAP transcriptional activity has been considered a gatekeeper of the immature fetal state ^61^. In cardiomyocytes, inhibition of YAP/TAZ induces a maturation gene program ^62^. Moreover, YAP deletion has been shown to promote premature senescence of mesenchymal stem cells ^63^. Hence, YAP/TAZ inactivation may be required for the maturation of multiple organs and tissues.

The identification of organ maturation checkpoints has several implications in regenerative medicine. Most adult somatic cells, such as cardiomyocytes and neurons, have limited regenerative capacity ^62,64^. In principle, activation of YAP/TAZ may push adult cells into immature states with regenerative potential. Indeed, activation of YAP in cardiomyocytes can promote heart regeneration in mice and pigs ^65,66^. However, as YAP/TAZ is regulated dynamically by upstream HPO1 and HPO2 signalings, custom-made strategies should be used to turn on YAP/TAZ activity in specific cells, and the duration of YAP/TAZ activation should also be strictly controlled to avoid tumorigenesis. Replacement of damaged tissues using *in vitro* expanded cells is another approach for tissue regeneration ^67^. However, *in vitro*-derived cells, including hepatocytes, are usually immature, affecting their functionality and imposing risk after transplantation ^68,69^. Activation of the Hippo pathway may promote cell maturation *in vitro*, and it would be interesting to test this idea experimentally in the future.

### The Hippo pathway in liver development and size control

The Hippo pathway has been considered an organ size regulator, as evidenced by rapid and reversible enlargement of livers upon YAP overexpression in mice ^14,18^. However, this concept has recently been challenged, mainly because organ size is not significantly changed in livers with *Yap/Taz* deletion ^21-24^, and hyperactivation of YAP/TAZ leads to ectopic growth, which does not recapitulate normal liver development ^22^. Hence, it is critical to review the role of the Hippo pathway in organ development and size control.

For organ size regulation, the function and shape of affected organs should be preserved, and it is ideal to have a proportional turnover of different parenchymal cells. The activation of YAP/TAZ in the liver results in dedifferentiation and transdifferentiation of hepatobiliary cells, and the hepatocytes to cholangiocytes ratio is inversely correlated with YAP/TAZ activity (Figs 3-5) ^25,70^. Complete inactivation of the Hippo pathway, such as deletion of *Lats1/2*, active YAP (YAP5SA) expression, and inactivation of both HPO1 and HPO2 (*Nf2^−/−^;Wwc^+/–^*or *Nf2^−/−^;Sav1^+/–^*), leads to massive accumulation of cholangiocytes (imCho1 and imCho2), ectopic liver growth, or rapid tumorigenesis (Fig 5) ^22,25,70^. In contrast, inactivation of HPO1, such as in *Sav1^−/−^;Wwc^−/−^*and *Mst1^−/−^;Mst2^−/−^* livers, results in accumulation of imCho1 and imHep and proportional liver enlargement; in these livers, the cholangiocytes to hepatocytes ratio is not dramatically perturbed, and the gene expression profile is similar to that of neonatal livers, indicating a relatively normal liver growth (Fig 3) ^15,16,19,20,25^. Hence, inactivations of the Hippo pathway can induce normal or ectopic liver growth, depending on YAP/TAZ activites.

The term Hippo pathway has been loosely defined and may refer to MST1/2 (Hippo), LATS1/2, or YAP/TAZ activity, which leads to ambiguity. Based on reported observations, HPO1 (the signaling from MST1/2) and HPO2 (the signaling from MAP4K1-7) may be considered canonical and noncanonical Hippo pathways, respectively. HPO1 is a predominant regulator of liver size, and its inactivation in the liver leads to intermediate YAP/TAZ activation and rapid liver enlargement ^15,16,19-21,25^. On the other hand, HPO2 plays a significant role in tumorigenesis, and indeed, *NF2* is the most frequently mutated Hippo pathway gene in human cancers ^17,25,71^. Concurrent HPO1 and HPO2 inactivation lead to YAP/TAZ hyperactivation, causing excessive transdifferentiation, tumorigenesis, and organ failure ^22,25,70^. Notably, LATS1/2 co-deletion or YAP5SA expression is rare even in pathological conditions, which may not represent a physiological approach to studying functions of the Hippo pathway in development.

The Hippo pathway has been shown to control organ size by regulating cell numbers. However, the upstream signal of the Hippo pathway in organ size control remains unknown ^14,18^. In livers, the number of hepatocytes should be the primary determinant of organ size, but the Hippo pathway should not be able to count cell numbers directly. Interstingly, both maturation and proliferation of hepatocytes are regulated by the HPO1. In a companion study, we have shown that YAP/TAZ directly regulates the expression of insulin-like growth factor II (IGF2), specifically in immature hepatocytes, and IGF2 is indispensable for liver enlargement following HPO1 inactivation ^72^. Hence, the maturation status of hepatocytes may be the mysterious upstream signal of HPO1. In postnatal developing livers, the maturation of hepatocytes will turn on HPO1, reduce IGF2 expression and cell proliferation, and restrict liver size (Fig 7B). It would be interesting to investigate if and how HPO1 senses the maturation of hepatocytes in the future.

## ACKNOWLEDGMENTS

This study is supported by grants from the National Key R&D Program of China (2020YFA0803202 and 2018YFA0800304), the National Natural Science Foundation of China (32425017, 32370770, and 32200570), the Science and Technology Commission of Shanghai Municipality (21S11905000), and the Shanghai Municipal Health Commission (2022XD049).

## EXPERIMENTAL MODEL AND SUBJECT DETAILS

### MOUSE MODELS

All mouse experiments in this study received approval from the Animal Ethics Committee of Shanghai Medical College, Fudan University, and were carried out under institutional guidelines. All mice utilized were of C57BL/6 backgrounds. *Wwc1*^F/F^, *Wwc2^F/F^*, and *Nf2^F/F^* were in-house generated as described in our prior work ^21,25^. *Sav1^F/F^*, *Yap^F/F^*, and *Taz^F/F^* mice were previously described ^24,66,73^. *Wwc1-LSL-mNeongreen* and *Nf2-LSL-mNeongreen* mice were in-house generated through CRISPR/Cas9-mediated homologous recombination. For these mice, the LoxP-STOP-LoxP-GS-Linker-mNeogreen expression cassette was knocked in at the termination codon site of the *Wwc1* or *Nf2* (isoform2) gene, as illustrated in Fig S4 and S5. Albumin-cre mice (Alb-cre, The Jackson Laboratory, #003574) were used for specific gene deletion in the liver. Ai9 (RCL-tdT) reporter mice (Ai9, The Jackson Laboratory, #007905), which express robust tdTomato upon Cre-mediated recombination, were used for lineage tracing. For hepatocyte-specific gene deletion using AAV8-TBG-cre (Obio Technology, #H5721), 1-month-old mice were injected with adeno-associated virus (1×10^11^ viral genomes for each mouse) via the lateral tail vein. Male and female mice were randomly assigned to experiments. In experiments involving tumorigenesis, mice were euthanized when severe abdominal enlargement or body weight loss (70% of control littermates) was observed. At the indicated age, mouse livers harvested were either fixed in formalin solution and paraffin-embedded or stored in liquid nitrogen for gene expression analysis. Some livers were directly embedded in OCT (Sakura), sectioned, and subjected to spatial transcriptomic analysis and some immunostaining.

For the construction of APAP-induced acute liver injury models, 2-month-old male mice were housed under a regular 12-hour light-dark cycle, with food removed 12 hours before APAP injection. APAP (dissolved in 0.9% saline, 300 mg/kg body weight) was administered intraperitoneally (i.p.). Food was then given to mice, and liver tissues were harvested at indicated time points (24, 48, or 96 hours after injection).

To establish DDC-induced liver injury models, 2-month-old male mice were fed with 0.1% DDC for the specified duration and were subsequently sacrificed after one month of continuous DDC exposure.

### METHOD DETAILS

#### Immunoblotting

Whole-cell lysates containing approximately 10 μg of proteins per sample were separated by sodium dodecyl sulfate-polyacrylamide gel electrophoresis (SDS-PAGE), and proteins were subsequently transferred onto nitrocellulose membranes. After blocking with 5% non-fat milk, the membranes were incubated with primary antibodies diluted in TBST (TBS containing 0.1% Tween-20) with 5% BSA overnight at 4°C, followed by incubation with secondary antibodies in 5% non-fat milk or BSA for 1 hr at room temperature. High-sig ECL Immunoblotting Substrate (Tanon, #180-501) was applied to the membranes, and chemiluminescence was detected using a Tanon 5200S imaging system.

#### Immunofluorescence staining

Paraffin-embedded liver tissues were sectioned (5 μm thickness), followed by rehydrated in a graded ethanol series (100% - 95% - 75% - 50% sequentially) and subjected to heat-induced antigen retrieval for 20 minutes using 10 mM sodium citrate buffer. Sections were pre-treated by blocking with 10% goat serum in PBST (PBS containing 0.1% Triton X-100) for 1 hr, followed by overnight incubation with primary antibodies diluted in PBST with 1% BSA at 4°C. Subsequently, tissue sections were incubated with secondary antibodies and DAPI (ThermoFisher, P36935) at room temperature for 1 hr after washing three times with PBST. Sections were then mounted using ProLong™ Gold Antifade Mountant. Images were captured with a Zeiss LSM900 confocal microscope or Olympus VS200 Slide Scanner.

#### Tyramide Signal Amplification (TSA) analysis

OCT-embedded liver tissues were sectioned (10 μm thickness), fixed in formalin solution, and treated with 3% H_2_O_2_ for 1 hr to quench endogenous peroxidase. After blocking with 1% BSA in PBST for 1 hr, sections were stained with anti-mNeongreen antibody (ChromoTek, 32F6, 1:200) in PBST overnight at 4°C. Following three washes, tissue sections were incubated with Envision anti-Rabbit/Mouse HRP antibody (DAKO, K5007) for 1 hr, and then stained by tyramide working solution (APExBIO, K1052, 1:100) for 10 mins. Sections were mounted, and images were captured with a Zeiss LSM900 confocal microscope.

#### Samples preparation and library construction

For single-cell RNA-sequencing samples, livers of 3-week-old mice were cut into pieces and rinsed using PBS to minimize blood contamination. The liver pieces were then enzymatically dissociated with dispase, collagenase type II and DNase I at 37°C for 20 mins for complete cell dissociation. The isolated single cells were immediately loaded onto a 10x Chromium Controller and then partitioned into nanoliter-scale Gel Beads-In-Emulsion (GEMs). Cells in GEMs were lysed, and the released RNA was immediately captured by barcoded beads, allowing for subsequent reverse transcription, amplification, fragmentation, adaptor ligation, and index PCR. Libraries were constructed using Chromium Single Cell 3’ Reagent Kits (V2 chemistry, 10x Genomics) according to the manufacturer’s instructions. For spatial transcriptomic samples, OCT-embedded frozen liver samples from 3-week-old mice that passed RNA quality control were processed for analysis. Samples were cut onto Visum slides, and the hematoxylin and eosin-staining images were exported as tiled tiffs for analysis. Samples were then further processed using the Visium Spatial Gene Expression Slide & Reagent Kit (10x Genomics) according to the Visium Spatial Gene Expression User Guide (CG000239, 10x Genomics) with an optimized permeabilization time of 15 mins.

#### Computational analysis

##### Single-cell RNA-seq data pre-processing and dimensionality reduction

Quality control and adapters trimming of scRNA-seq fastq data are conducted using the fastp pipeline ^74^. The single-cell data is further mapped with a standard cellranger^75^ (Version 7.1.0) pipeline to mm10 reference pre-built by 10X Genomics, which is consistent with single-cell data. The Seurat^76^ (version 4.0.6) package was used for data processing. The outliers were identified and removed based on library size, number of expressed genes, and mitochondrial genes determined individually for each sample. Doublets were identified and excluded using the DoubletFinder^77^ (v2.0.3) package in R. Briefly, artificial doublets were simulated by merging pairs of cells, and each cell was assigned a probability score reflecting its likelihood of being a doublet. The threshold for doublet removal was set based on the expected doublet rate, calculated as a proportion of total cells (7.5% doublet formation rate in this study). The cleaned dataset is merged and further processed sequentially for normalization, scale, finding variable features, and PCA calculation. Dimensionality reduction and unsupervised clustering are performed using Seurat RunUMAP function loading with 30 PCs. To find clusters, an SNN graph-based clustering algorithm with the same PCs used for RunUMAP was imported into FindClusters.

##### Visium data pre-processing

Quality control and adapters trimming of scRNA-seq fastq data are conducted using the fastp pipeline ^74^. The bright field image of spatial transcriptomics data is processed with Loupe Browser (Version 6.4.0). After manually aligning the bright field image and removing the spots outside the tissue-captured area, the results were exported as json files. The json files and quality-controlled reads files are processed with a standard spaceranger (Version 2.0.0) pipeline to 10x distributed mm10 reference. Main computational analysis of spatial read-count matrices was performed using Seurat (version 4.0.6) package in R (v4.0.2) or Scanpy (v1.10.2) in Python (3.9). Similar to scRNA-seq preprocessing, outliers of Visium spots are determined by 3 metrics, including library size, number of expressed genes, and mitochondrial genes calculated individually for each sample. The cleaned data is further processed sequentially for normalization, scaling, and calculation of highly variable genes (HVG).

##### Data integration

To integrate multiple Visium datasets or scRNA-seq data from different sources while preserving biological variability and mitigating batch effects, we employed the Reciprocal Principal Component Analysis (RPCA) method implemented in the Seurat package. A common set of features was identified across all datasets using the SelectIntegrationFeatures function, specifying nfeatures = 2,000 to ensure a robust set of genes for integration. We identified integration anchors using the FindIntegrationAnchors function. Data from different livers were then integrated based on common anchors. PCA and UMAP were calculated based on the integrated assay according to Seurat standard spatial workflow. To determine the clusters of Spatial spots in an unsupervised manner, the Louvain algorithm for community detection using the FindClusters function were applied, which clusters cells by optimizing the modularity of the SNN graph.

##### Evaluation of Integration Performance

To assess the effectiveness of the data integration, we implemented a comprehensive evaluation strategy that examines both the correction of batch effects and the preservation of biological variability. This evaluation involved quantitative metrics, visualization techniques, and statistical analyses to ensure that the integrated dataset accurately reflects true biological signals while minimizing technical artifacts. We calculated the Batch LISI (bLISI) and Cell Type LISI (cLISI) using the LISI package^78^ to quantify batch mixing and cell type separation in the integrated data.

##### Differential gene expression analysis

Differential gene expression analysis was conducted using Seurat’s FindAllMarkers function to detect genes uniquely expressed in each cluster. The analysis employed the Wilcoxon rank-sum test, a non-parametric method suited for comparing two groups, which was applied to each cluster to find differentially expressed genes (DEGs). We set an initial threshold, requiring genes to have a log fold-change (logFC) above 0.25 and be expressed in at least 10% of the spots within a cluster to be considered significant. Further filtering was done to refine the selection, isolating marker genes with a logFC greater than 0.5 and an adjusted p-value below 0.05.

##### Gene Ontology (GO) enrichment analysis

To explore the biological functions associated with the differentially expressed genes (DEGs), we performed Gene Ontology (GO) enrichment analysis using the clusterProfiler package (version 4.12.6) in R. For the GO annotations, we used the org.Mm.eg.db annotation package (version 4.4), which provides the mapping between mouse gene identifiers and their corresponding GO terms. To control for multiple hypothesis testing, p-values were adjusted using the Benjamini-Hochberg method. GO terms with an adjusted p-value < 0.05 were considered statistically significant.

##### Monocle 3

To infer the transition and continuity from the sequencing data, we used highly variable genes and applied learnGraph, which learns the structure of the dataset manifold as developmental trajectory and ordered cells by calculating a pseudo-time-assisted trajectory from selected root cells. For branch pruning, we adopt a heuristic pruning method aimed at refining the principal graph by excising diminutive branches^79^. To retrieve the differential expressed genes, we use a Graph-autocorrelation analysis pipeline within monocle3 to evaluate genes that vary over a trajectory.

##### Partition-based Graph Abstraction (PAGA)

The PAGA algorithm maps coarse-grained connectivity structures of complex manifolds^38^. By quantifying the connectivity of partitions (groups, clusters) of the single-cell graph, partition-based graph abstraction, a graph is generated in which edge weight represents the confidence in the presence of connections. An additional denoising step was taken to calculate distances within a few diffusion components, as described by Schiebinger et al.^80^.

##### Gene regulator network analysis

To identify upstream regulators important for the development of cholangiocytes and hepatocytes, we have performed SCENIC (Single Cell Regulatory Network Inference and Clustering) with a segregated raw expression matrix for the cell type of interest. For quality control, we kept cells with at least 200 genes expressed and only genes that were expressed in at least 3 cells, and doublets were identified and filtered out using DoubletFinder ^77^. SCENIC is then performed with steps involving co-expression analysis, Motif discovery, and cell scoring in a sequential manner to identify and score regulons based on the expression of their regulated target genes. A gamma distribution is performed to fit the regulon score and identify significant differential transcription factors (TFs).

##### Gene set scoring

To quantify the activity of specific gene sets and assess their expression patterns across different cell populations, we implemented a gene expression scoring method using the Seurat package. We curated gene sets representing specific biological pathways or cellular functions of interest. These gene sets were obtained from reputable databases such as the Molecular Signatures Database (MSigDB) or compiled from relevant literature. To ensure robust scoring, genes not detected in our dataset were removed from the gene sets. Only genes expressed in at least 1% of the cells were retained to avoid skewing the scores due to low or absent expression. We utilized Seurat’s AddModuleScore function to compute module scores for each cell based on the expression of genes within the predefined gene sets.

##### Modeling of Spatial relation

To capture the spatial architecture of the tissue, we represented the Visium slide locations (spots) as nodes in a graph, with edges denoting neighborhood relationships between adjacent spots. Each spot’s neighbors were defined by the number of surrounding hexagonal rings, which allowed us to systematically control and define the spatial proximity of each node. Using this information, we constructed a spatial graph that models the local environment of each spot, effectively capturing how regions of interest are spatially organized across the tissue. This graph-based approach enables a quantitative understanding of spatial proximity and allows for a more nuanced analysis of zone-specific gene expression patterns, facilitating the identification of spatially co-regulated gene networks and distinct tissue microenvironments.

##### Cell2location

To resolve fine-grained spatial distributions of cell types, we utilized Cell2location ^81^, a Bayesian modeling framework designed to deconvolve cell type abundances from mini-bulk spatial transcriptomic data. Cell2location effectively decomposes multi-cell spatial transcriptomics data into spatially resolved estimates of cell-type abundance. The scRNA-seq reference map was created by merging our in-house scRNA-seq data with publicly available time-series mouse liver scRNA-seq data, ensuring comprehensive cell-type representation ^82^. To mitigate batch effects between the scRNA-seq and spatial datasets, we employed a gene-specific scaling parameter that accounts for technology-dependent differences in sensitivity. Expression signatures for each cell type were derived by calculating the average expression counts of each gene across all cells of that type, ffiltering genes expressed in at least three cells. To obtain spatially resolved cell type abundances, we applied a hierarchical non-negative matrix decomposition of the gene expression profiles at spatial locations (i.e., Visium spots containing multiple cells) into reference signatures. Each Visium section was analyzed independently with the following modified parameters: n_iter: 30,000; n_samples:1,000;cell_per_spot: 10; factors_per_spot: 4; The deep learning model for the reference data was trained on a Nvidia RTX3080 GPU with CUDA version 11.2, using 10,000 to 30,000 epochs depending on the convergence of the loss function. Finally, we visualized the absolute mRNA contribution of each cell population to each spatial spot by analyzing the 5th percentile of the posterior distribution for the number of mRNA molecules confidently assigned to each cell type.

##### Shannon Entropy Calculation

Cell abundances is calculated as described in Method Cell2loation and normalization is performed as following:

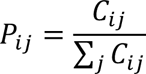

Where *P_ij_* is the normalized proportion of cell type *j* in spot *i*, and *C*_*ij*_ is the abundances of cell type *j* in spot *i*.

For each spot *i*, calculate the Shannon Entropy as following:

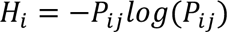

Where is *H_i_* the Shannon Entropy for spot *i*.

##### Quantification and statistical analysis

We primarily used Kruskal-Wallis tests for statistical differences, incorporating the Benjamini-Hochberg False Discovery Rate (FDR) for multiple hypothesis adjustments. Spearman correlations assessed relationships, and gene expression fold changes between cell groups were calculated using log2(ratio + minimal positive value) to avoid division by zero.

## Data availability

The sequencing data has been deposited to The National Genomics Data Center (https://ngdc.cncb.ac.cn) with a project number of PRJCA025196.

---

**Figure S1.**
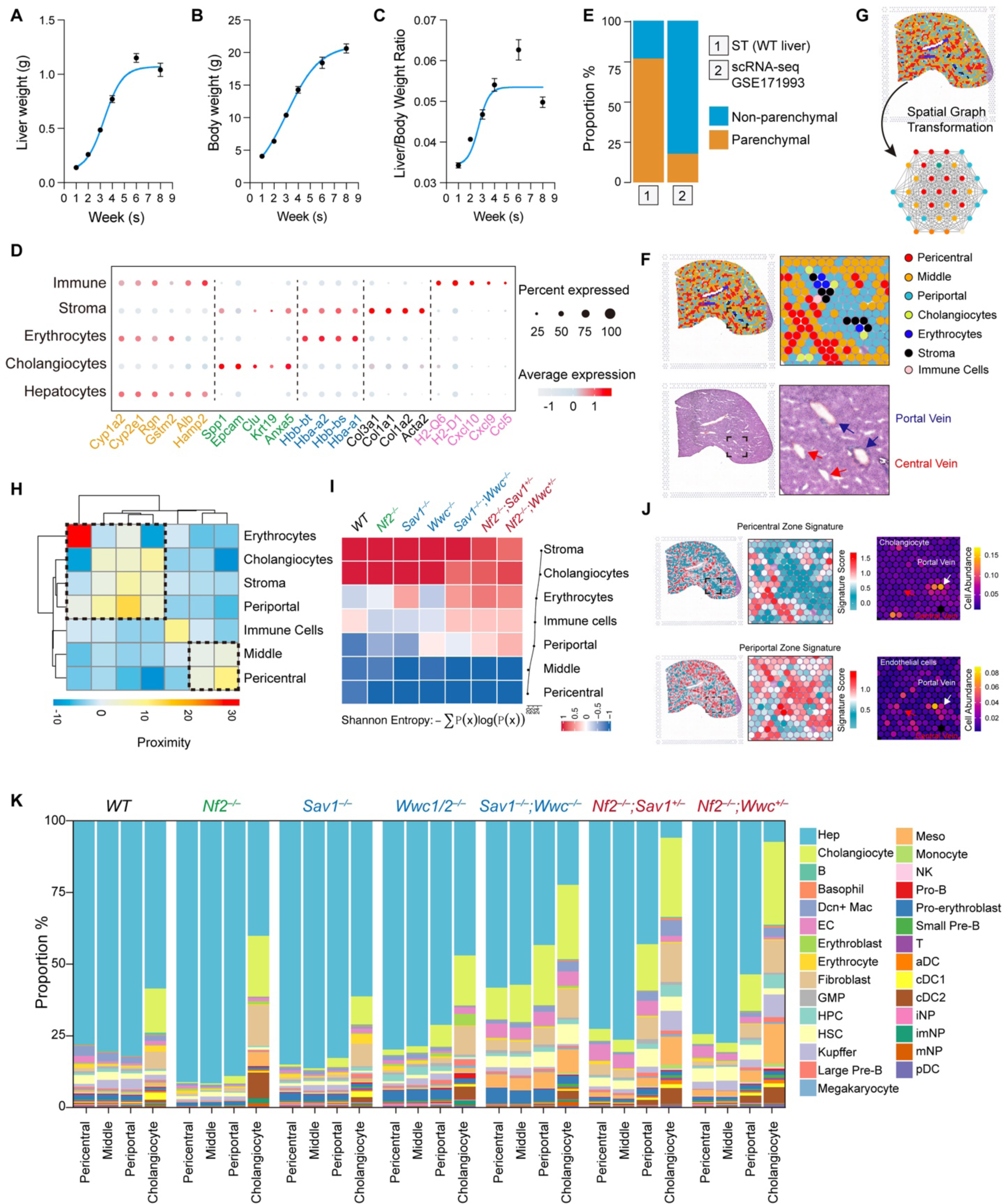
Perturbations of the Hippo pathway result in dynamic changes in cellular compositions in the liver. (A-C) Postnatal growth of mouse livers. Mouse body weight (A), liver weight (B), and body liver-to-weight (C) during postnatal development are shown. (D) Dot plot of marker gene expression. The dot size represents the percentage of spatial spots with non-zero expression, and the color indicates the average expression value within each spatial spot. (E) Proportion of hepatobiliary cells in ST and scRNA-seq datasets. (F) Molecular features align with histological structures. Top: Spatial spots defined by transcriptomics fingerprint. Bottom: H&E staining images. (G) Figure illustrating the spatial transformation of spatial spots. The location of each spot is modeled by a hexagonal grid. (H) Proximity heatmap of spatial spots. The heatmap illustrates spatial proximity between spots. (I) Heatmap showing Shannon entropy across different spots in various samples. The hepatocyte spots exhibit the lowest entropy across all genotypes. (J) liver zonation revealed from ST data. Left: zonation assigned to each spatial spot. Right: spatial projection of decomposed cell abundances. Ductular cells are specifically localized near the portal vein, while endothelial cells exhibit a peak distribution around the portal (white arrow) and central vein (red arrow). (K) Cell composition in different liver samples. Cell abundances in each spatial spot are calculated using a reference scRNA-seq data of mouse liver (GSE171993). The proportion of hepatocyte (Hep), cholangiocyte, endothelial cell (EC), hepatic stellate cell (HSC), fibroblast, mesothelial cell (Meso), megakaryocyte, erythroid cell (pro-erythro-blast, erythroblast, and erythrocyte), T cell, natural killer (NK) cell, B cell (Pro-B, large pre-B, small pre-B, and B), dendritic cell (classical dendritic cell 1 – cDC1, classical dendritic cell 2 – cDC2, plasmacytoid dendritic cell – pDC, and activating dendritic cell – aDC), monocyte, Dcn+ macrophage (Dcn+ Mac), Kupffer cell,neutrophils (immature neutrophil – iNP, intermediate mature neutrophil – imNP, and mature neutrophil – mNP), basophil, granulocyte-monocyte progenitor (GMP), and hematopoietic progenitor cell (HPC) in particular zones and genotypes is shown.

**Figure S2.**
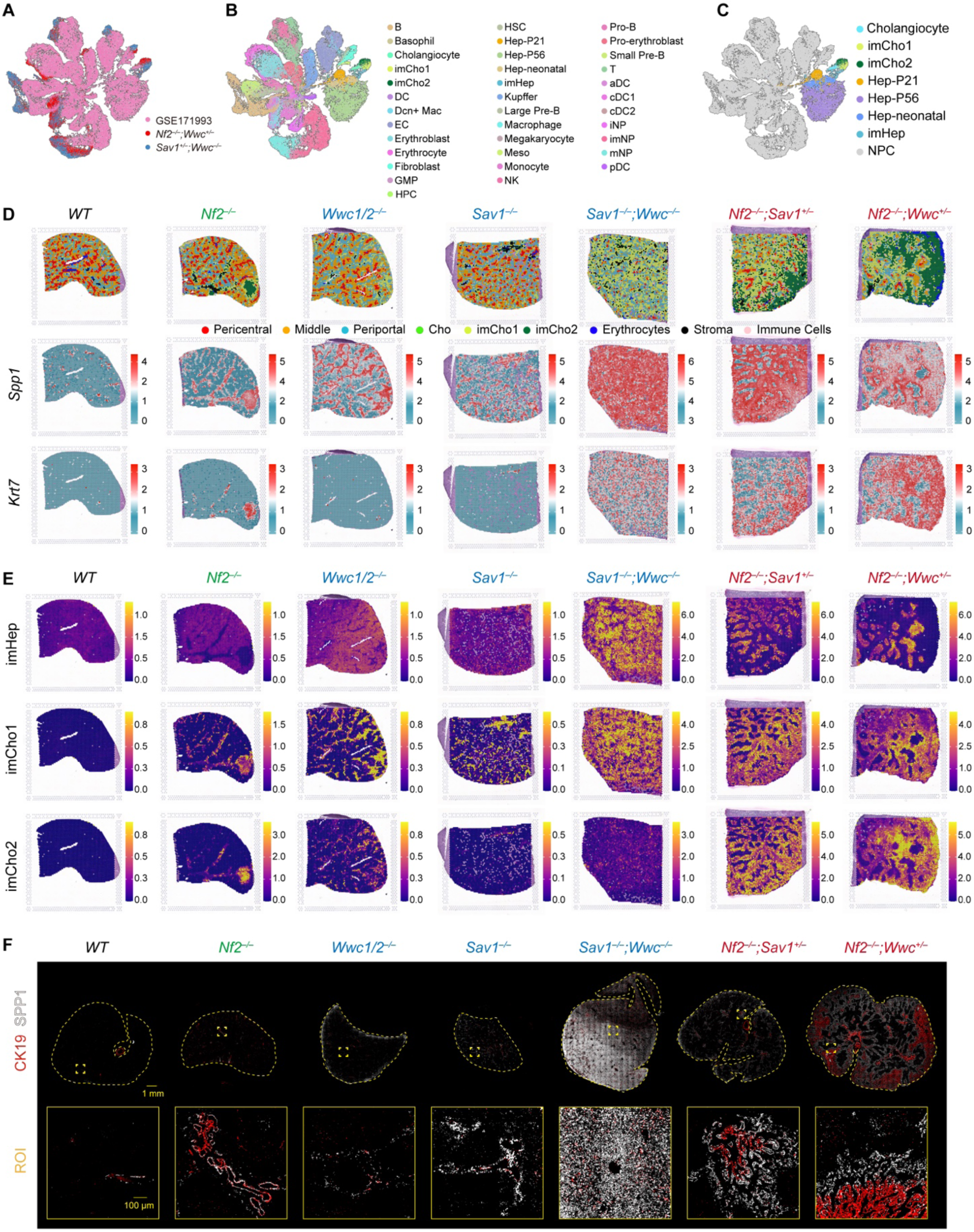
Spatial distribution of three pools of immature hepatobiliary cells. (A-C) UMAP representation of the integrated liver cell atlas. Public scRNA–seq data (GSE171993) and scRNA–seq data from *Nf2^−/−^;Wwc^+/–^* and *Sav1^−/−^;Wwc^−/−^* mouse livers are intergrated. Cells are colored by batch (A) or cell type (B). Hepatocytes and cholangiocytes are highlighted (C). (D) Spatial expression of *Spp1* and *Krt7* in different livers. (E) Distribution of imCho1, imCho2, and imHep in different livers. (F) Expression of CK19 and SPP1 in different livers. Liver sections are co-stained for CK19 (red) and SPP1 (white). CK19^high^SPP1^low^ and CK19^low^SPP1^high^ cells indicate imCho2 and imCho1 cells, respectively. Large scanned images and ROI with higher magnifications are shown. Scale bars: 1 mm or 100 μm as indicated.

**Figure S3.**
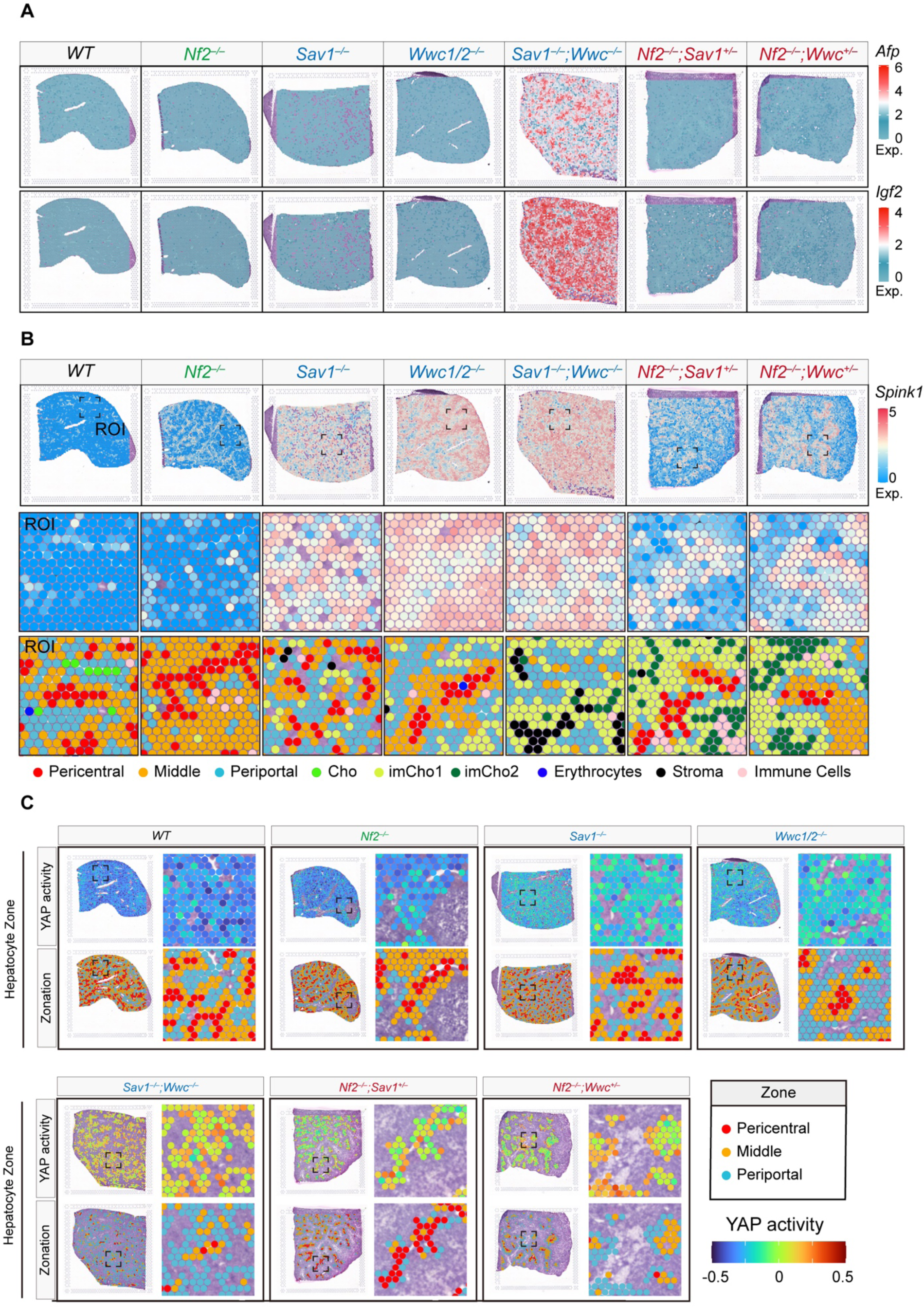
HPO1 inactivation leads to the expansion of immature hepatocytes. (A) Spatial expression of hepatoblast markers (*Afp* and *Igf2*) in different livers. (B) High expression of *Spink1* in HPO1 defective livers. (C) Spatial map of YAP activity in hepatocyte regions. Each spatial spot is colored according to the calculated YAP activity, and corresponding zonations of hepatocytes are shown.

**Figure S4.**
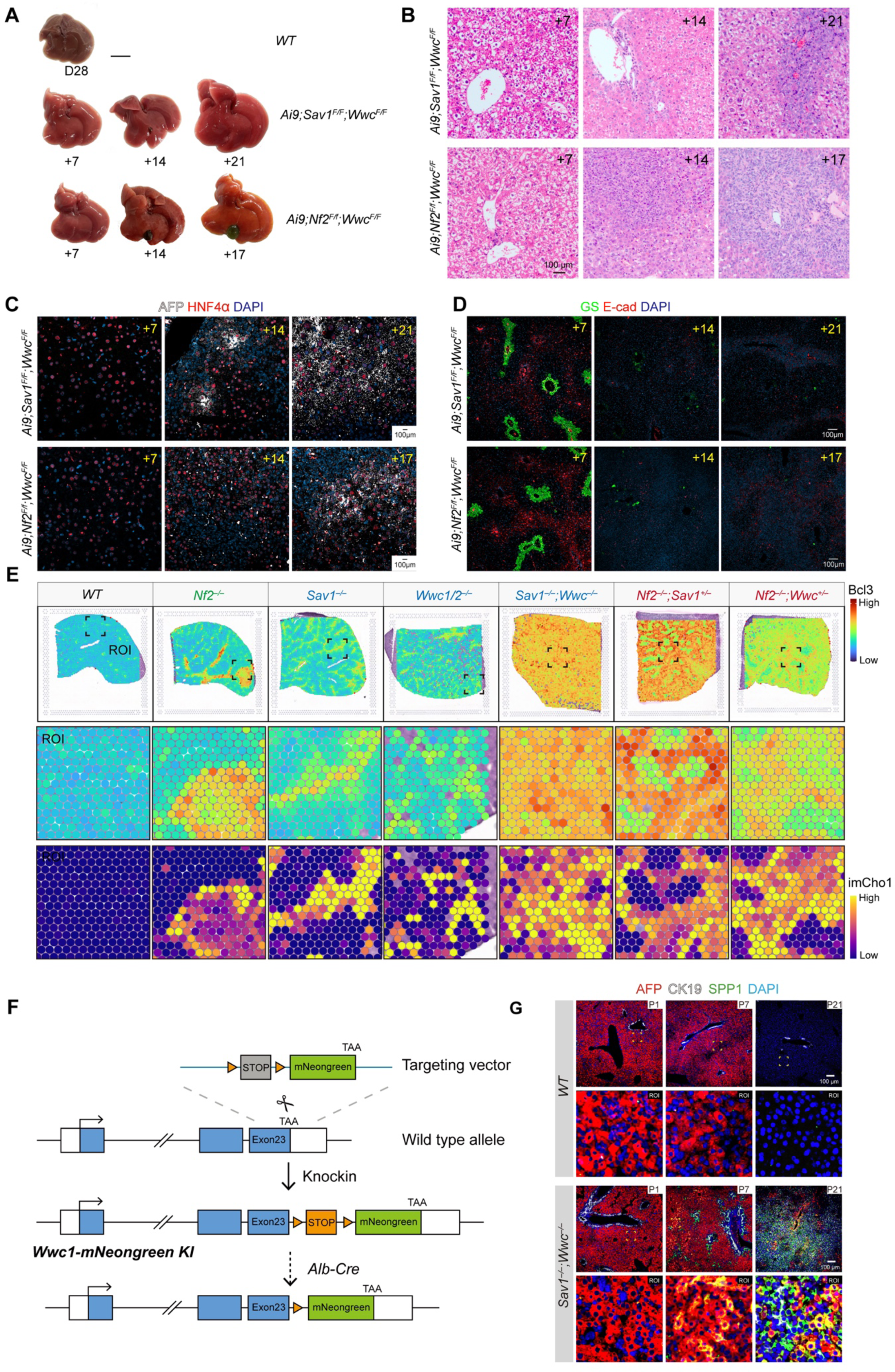
Defective Hippo signaling induces hepatocyte reprogramming and transdifferentiation into imCho1. (A and B) Gross images (A) and H&E staining (B) of mouse livers at different time points after AAV-TBG-Cre tail vein injection. At 17 or 21 days post-injection, the size of livers is robustly enlarged, and bile duct expansion is evident. Scale bars: 1 cm (A) or 100 μm (B). (C and D) Immunostaining of liver sections. C: AFP (white) and HNF4α (red). D: GS (green) and Ecad (red). The same livers are used in A and B. Scale bar: 100 μm. (E) Bcl3 activity calculated in different livers. Bcl3 activity is calculated based on the expression of Bcl3 target genes and projected onto the spatial transcriptomic matrix. The distribution of imCho1 cells in corresponding regions is also shown. (F) Schematic diagram illustrating the strategy for generating *Wwc1*-LSL-mNeonGreen knock-in mice. LSL (loxP-STOP-loxP) was removed upon liver-specific Alb-Cre mediated recombination. (G) Immunostaining of liver sections. AFP (red), CK19 (white), and SPP1 (green) are stained for liver sections from WT or *Sav1^−/−^*;*Wwc^−/−^* mouse (*Alb*-cre) livers at indicated ages. Scale bar: 100 μm.

**Figure S5.**
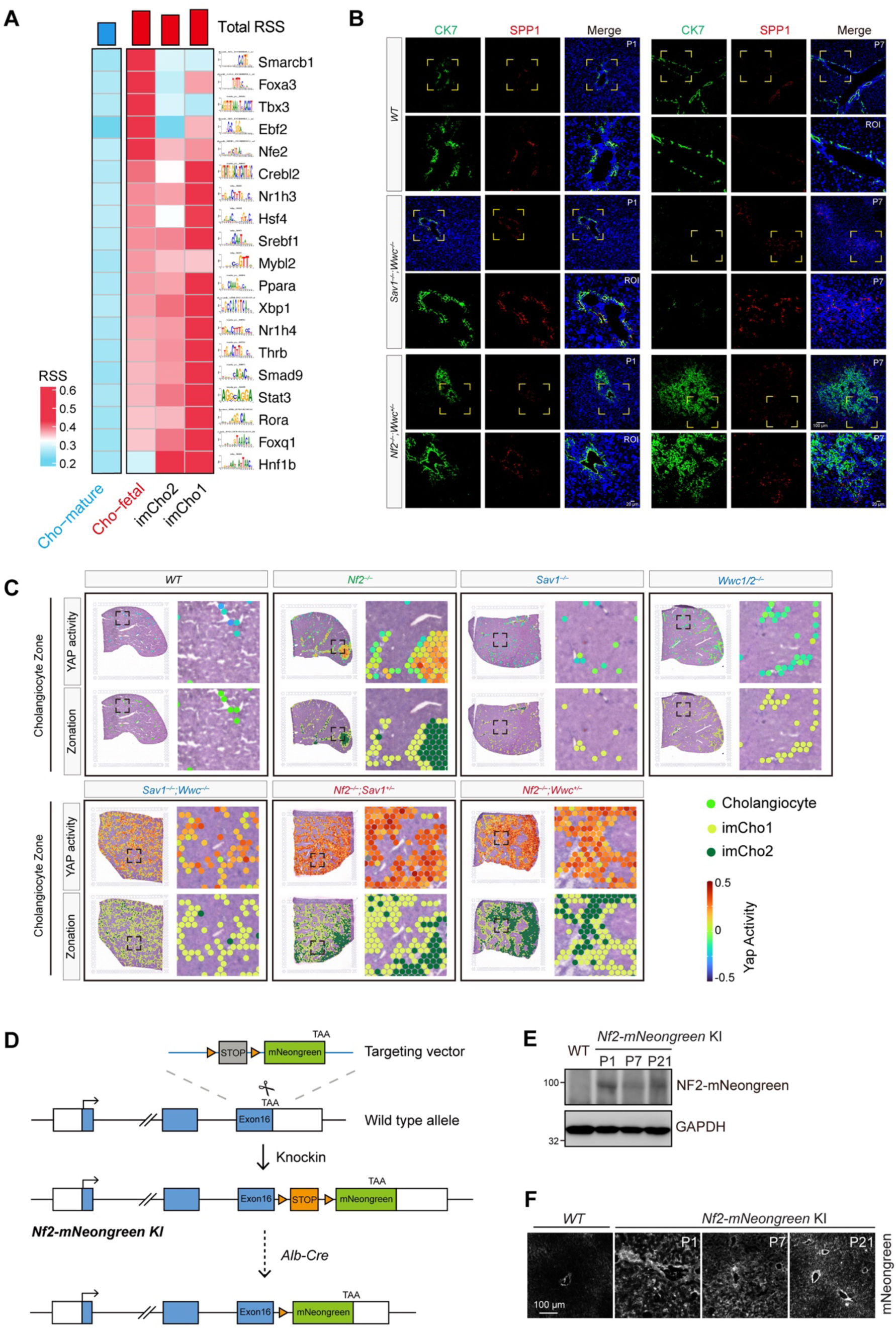
Inactivation of HPO2 leads to the accumulation of imCho2 cells. (A) Heatmap showing Regulon Specific Scores (RSS) calculated by SCENIC analysis. imCho1, imCho2, and immature fetal cholangiocytes exhibit similar transcription factor activities. (B) Immunostaining of CK7 (green) and SPP1 (red). Liver sections from wild type (WT), *Sav1^−/−^;Wwc^−/−^* mice, and *Nf2^−/−^;Wwc^+/–^* mice at P1 and P7 after birth are analyzed. Scale bars: 100 μm or 20 μm as indicated. (C) Spatial projection map of YAP/TAZ activity in cholangiocyte regions. (D) Schematic diagram illustrating the strategy for generating *Nf2*-LSL-mNeonGreen knock-in mice. LSL (loxP-STOP-loxP) is removed upon liver-specific *Alb*-Cre-mediated recombination. (E and F) Temporal analysis of *Nf2*-mNeonGreen expression. Liver samples were obtained from wild type (WT) mice and *Nf2*-mNeonGreen knock-in mice on postnatal days P1, P7, or P21. Protein expression (E) and immunostaining (with TSA) analysis (F) are shown.

**Figure S6.**
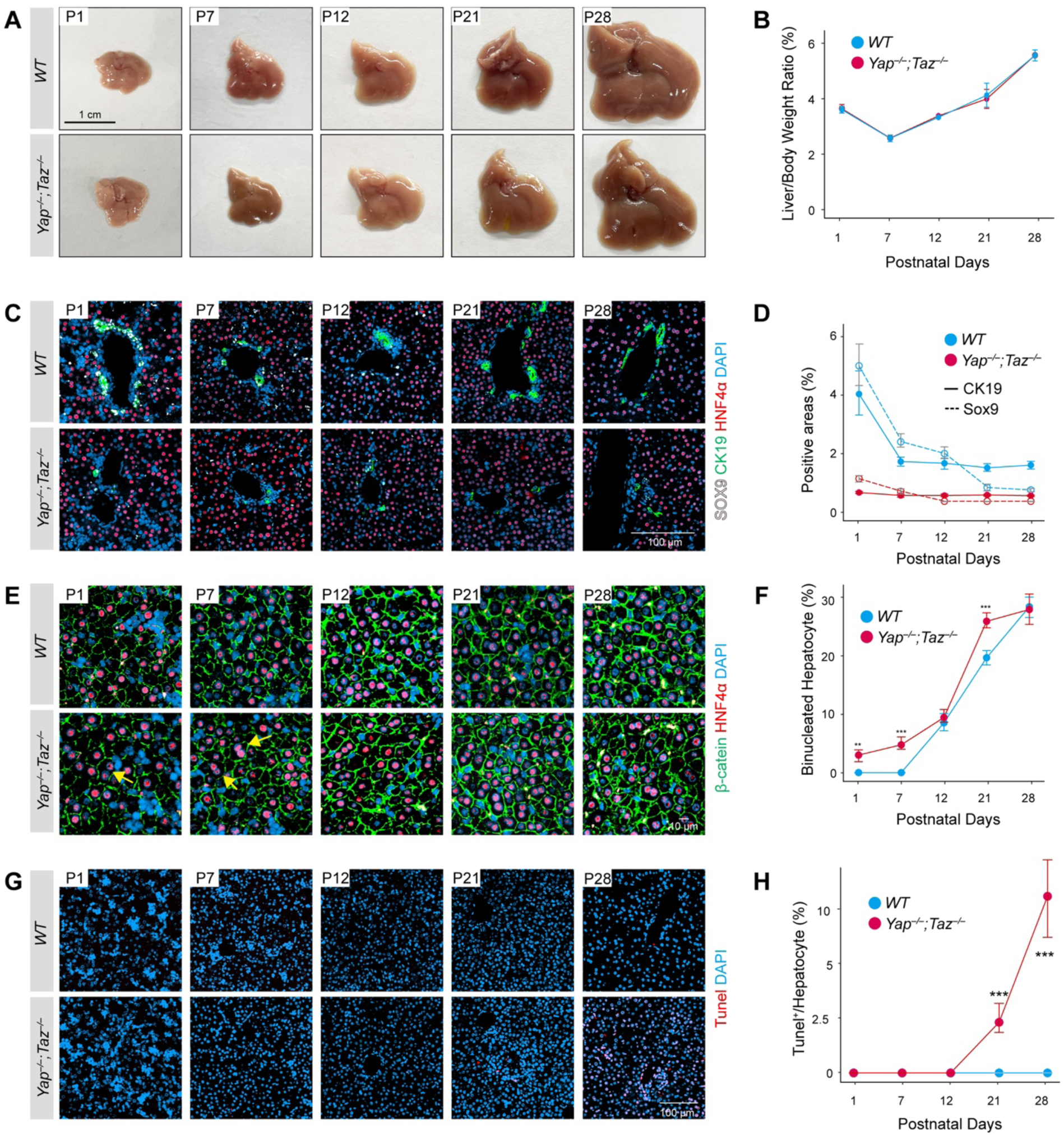
*Yap/Taz* deletion promotes liver maturation and tissue damage. (A and B) Liver-specific *Yap/Taz* deletion has a marginal effect on liver size. Gross liver morphologies in wild type (WT) and *Yap^−/−^;Taz^−/−^* mice are shown at postnatal days (P) 1, 7, 12, 21, and 28 after birth. Scale bar: 1 cm. Statistical analysis is shown in B. (C and D) Immunostaining of Sox9, CK19, and HNF4α. Tissue sections are from livers described in A. Statistical analysis is shown in D. Scale bar: 100 μm. (E and F) Immunostaining of β-catenin and HNF4α. Tissue sections are from livers described in A. Quantification of binucleated hepatocytes is shown in F. Immunofluorescence analysis reveals altered β-catenin and HNF4α expression in *Yap/Taz* dKO livers. Liver sections from wild type (WT) and liver-specific *Yap/Taz* double knockout (KO) mice were co-stained for β-catenin (a marker of Wnt signaling) and HNF4α (hepatocyte marker). Scale bar: 100 μm. (G and H) TUNEL staining. TUNEL-positivity (red) indicates dying cells. Scale bar: 100 μm. Quantification is shown in H. Data are presented as mean ± SEM from at least three independent biological replicates.

**Figure S7.**
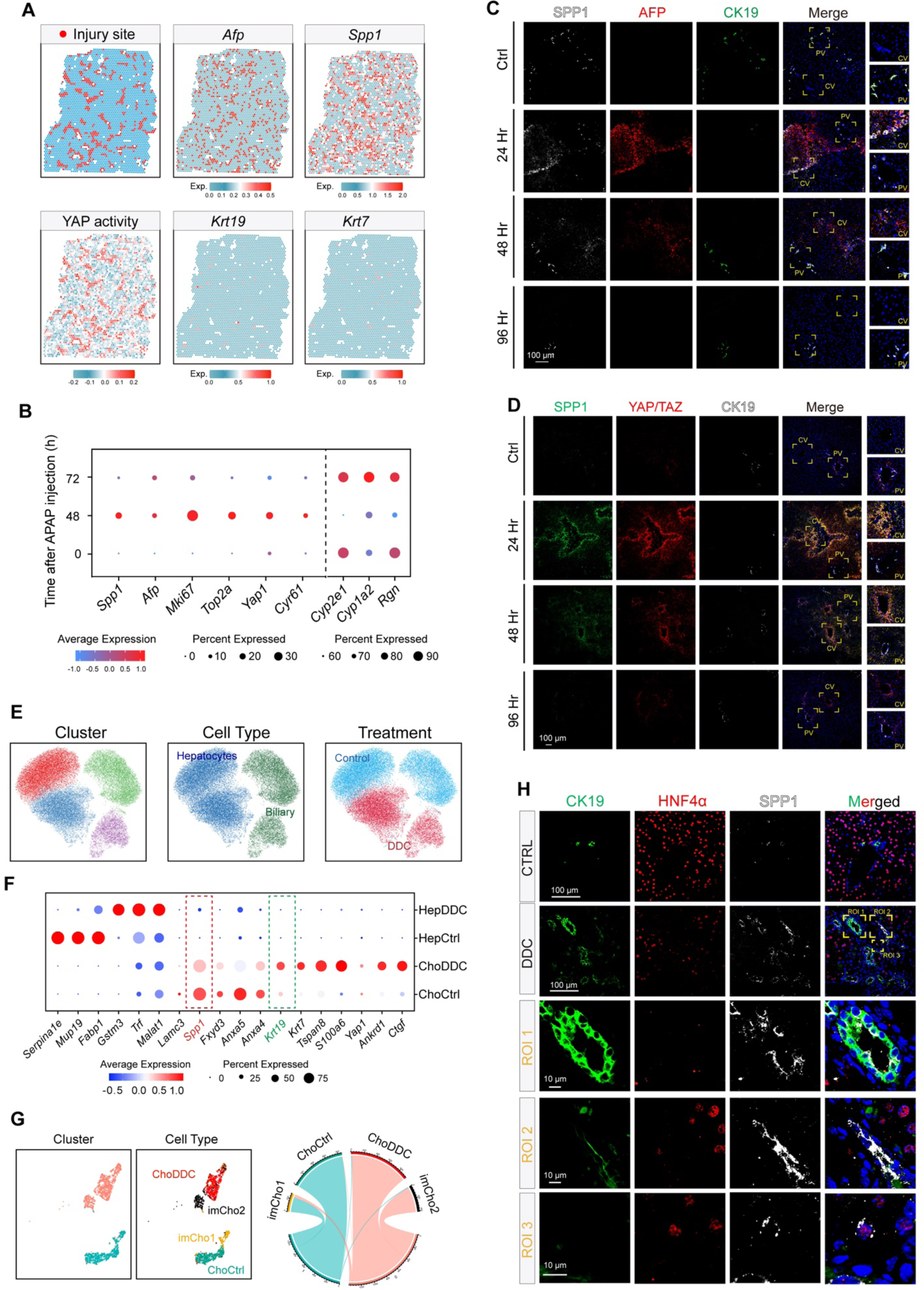
Liver damage induces patterned YAP/TAZ activation and the accumulation of immature cells. (A) Spatial analysis shows YAP activation and imHep signature in livers with APAP-induced injury. 48 hours after APAP injection, liver sections from mice were assessed for injury site distribution and expression of imHep markers (*Afp*, *Spp1*), cholangiocyte markers (*Krt19*, *Krt7*), and YAP activity. (B) scRNA-seq reveals dynamic changes in hepatocyte gene expression after APAP treatment. Dot plots display the average expression of *Afp*, *Spp1*, *Krt19,* and additional genes in hepatocytes from the liver at different time points following APAP injection. (C) Immunostaining of SPP1, AFP, and CK19. Liver sections from control mice and mice at 24, 48, and 96 hours after APAP injection were analyzed. The dashed yellow box highlights a region of interest (ROI) for magnified visualization. Hepatocytes at the edge of the injury site transition to imHep at 24 and 48 hours post-APAP injection, returning to normal 96 hours later when liver regeneration is complete. Scale bar: 100 μm. (D) Immunostaining of SPP1, YAP/TAZ, and CK19. The same samples in C are analyzed. Scale bar: 100 μm. (E) UMAP representation of integrated hepatobiliary cell atlas under normal conditions and after DDC treatment. (F) Dot plot showing expression of selected genes. The scRNA-seq data of hepatocytes and cholangiocytes under control (Ctrl) and DDC-treated conditions are analyzed. (G) UMAP representation of integrated cholangiocyte cell atlas by integrating cholangiocyte from norm and DDC condition to imCho1 and imCho2. On the right panel, a Circos diagram indicates the proportional distribution of integrated single-cell data. (H) Immunostaining of HNF4α, SPP1, and CK19 expression in livers with DDC-induced liver injury. Scale bar: 100 μm.

